# Ecological divergence and hybridization of Neotropical *Leishmania* parasites

**DOI:** 10.1101/824912

**Authors:** Frederik Van den Broeck, Nicholas J. Savill, Hideo Imamura, Mandy Sanders, Ilse Maes, Sinclair Cooper, David Mateus, Marlene Jara, Vanessa Adaui, Jorge Arevalo, Alejandro Llanos-Cuentas, Lineth Garcia, Elisa Cupolillo, Michael Miles, Matthew Berriman, Achim Schnaufer, James A. Cotton, Jean-Claude Dujardin

## Abstract

The tropical Andes is an important natural laboratory to understand speciation and diversification in many taxa. Here, we examined the evolutionary history of parasites of the *Leishmania braziliensis* species complex based on whole genome sequencing of 67 isolates from 47 localities in Peru. We firstly show the origin of near-clonal Andean *Leishmania* lineages that diverged from admixed Amazonian ancestors, accompanied by a significant reduction in genome diversity and large structural variations implicated in host-parasite interactions. Beside a clear dichotomy between Andean and Amazonian species, patterns of population structure were strongly associated with biogeographical origin. Molecular clock analyses and ecological niche modeling suggested that the history of diversification of the Andean lineages is limited to the Late Pleistocene and intimately associated with habitat contractions driven by climate change. These results support a wider model on trypanosomatid evolution where major parasite lineages emerge through ecological fitting. Second, genome-scale analyses provided evidence of meiotic recombination between Andean and Amazonian *Leishmania* species, resulting in full-genome hybrids. The mitochondrial genome of these hybrids consisted of homogeneous uniparental maxicircles, but minicircles originated from both parental species, leaving a mosaic ancestry of minicircle-encoded guide RNA genes. We further show that mitochondrial minicircles - but not maxicircles - show a similar evolutionary pattern as the nuclear genome, suggesting that biparental inheritance of minicircles is universal and may be important to alleviate maxicircle-nuclear incompatibilities. By comparing full nuclear and mitochondrial genome ancestries, our data expands our appreciation on the genetic consequences of diversification and hybridization in parasitic protozoa.

## INTRODUCTION

Exploring natural genetic variation has been instrumental in understanding how and under which circumstances new species originate. South America encompasses a large fraction of the global biodiversity, representing one of the most species diverse hotspots on Earth. This is partly because diversification of Neotropical taxa has been influenced by a rich geological and climatic history, including large-scale reconfigurations of the landscape through the Neogene uplift of the Andes^1^ and habitat instability through Pleistocene climatic cycling^2^. In particular the Andes – due to a complex interplay of history, geography and ecology – represents an epicenter for species diversification in birds, reptiles, insects and plants^3–8^. However, little is known about the role of the Andes shaping the evolution of parasitic micro-organisms.

Here, we used whole-genome sequencing to study the evolutionary history of parasites of the *Leishmania braziliensis* species complex in Peru, one of the biologically richest and most diverse regions on Earth. The lowland species *L. braziliensis* is a zoonotic parasite circulating in a diverse range of wild mammals^9^ in Neotropical rainforests. It is one of the major causes of cutaneous leishmaniasis in Latin America, and also causes a relatively high frequency of severe mucocutaneous disease (known locally as *espundia*) where the parasite spreads to mucosal tissue. Human infections appear to be a spillover from this sylvatic life cycle, and are probably not important in transmission. In contrast, the montane species *L. peruviana* is largely endemic to the Pacific slopes of the Peruvian Andes. It is transmitted exclusively in peri-domestic xerophytic environments and causes a disease of altitude (1300-2800 m), known locally as *uta*, a benign form of cutaneous leishmaniasis. Both *Leishmania* species were initially shown to be different at only one of 16 enzymatic loci^10^, but subsequent molecular analyses showed that they correspond to two distinct monophyletic clades of the *Viannia* subgenus^11–13^. Molecular karyotyping further revealed that *L. braziliensis* was karyotypically homogeneous in contrast to *L. peruviana* that was genetically subdivided into different biogeographical regions across and along the Peruvian Andes^14,15^. These species differences persist despite the report of a number of parasites presenting hybrid marker profiles^16,17^.

Our understanding of the genetics of diversification and hybridization in *Leishmania* is increasingly informed by the genomic revolution. Whole genome sequencing data of hundreds of isolates revealed the global^18^ and local^19,20^ genome diversity of the Old-World *L. donovani* species complex, although genome studies on New-World *Leishmania* species remain scarce and limited to a few isolates^21–23^. Genome-scale analyses also provided strong evidence for genome-wide patterns of recombination in presumed clonal *Leishmania* species^24–26^, including evidence of classical crossing over at meiosis^24^. While these studies revolutionized our understanding of the fundamental biology of parasitic protozoa, information on the structure, diversity and evolution of their mitochondrial genome remains fragmentary. This is mainly because of the extraordinary complexity of the mitochondrial DNA of trypanosomatids, consisting of a giant network of thousands of heterogeneous minicircles (0.5-2.5 kb in size, depending on the species) interlaced with 20-50 homogeneous maxicircles (20-30 kb)^27^. The minicircles encode guide RNA (gRNA) genes that are responsible for directing an elaborate U-indel RNA editing process that generates translatable maxicircle-encoded transcripts^27^. This process of RNA editing in the kinetoplast is a unique biological characteristic shared by all *Trypanosomatidae*, including other species of medical and veterinary importance such as *Trypanosoma brucei*, *T. vivax* and *T. congolense*. Correct RNA editing is essential for parasite viability^28^ and depends on a functionally complete set of minicircles^29^. However, little is known about the natural variation of minicircle complexity.

Here, we present the first study that examines the diversification and hybridization of parasitic protozoa based on a joint analysis of complete nuclear and mitochondrial genomes. After mapping sequences against a PacBio assembly including 35 chromosomes and a complete maxicircle, unaligned reads were used to assemble, circularize and annotate full sets of mitochondrial minicircles. We show that patterns of population genomic structure were strongly associated with biogeographical origin, and suggest that speciation was driven by ecological re-arrangements during late Pleistocene climatic cycling. In addition, we demonstrate that interspecific meiotic recombination resulted in uniparental inheritance of maxicircles but biparental inheritance of minicircles, leaving a mosaic ancestry of minicircle-encoded guide RNA genes. We discuss the potential role of biparental inheritance of minicircles in preserving minicircle complexity and mito-nuclear compatibility in trypanosomatid parasites.

## RESULTS

### Natural variation of the *L. braziliensis* species complex

The genomes of 31 *L. peruviana*, 23 *L. braziliensis* and 13 hybrid *L. braziliensis* × *L. peruviana* isolates (Supp. Fig. 1; Supp. Table 1) were sequenced at a median 70x depth (mean = 73x; min = 40x; max = 123x). For comparative purposes we also included sequencing data of one isolate from the closely related *L. panamensis* species (Supp. Table 1). All sequence data were aligned against a novel long-read assembly of *L. braziliensis* M2904, and for every isolate we determined the accessible genome that is characterized by sufficient mapping quality, base quality and read depth. This revealed that 90.9-92.2% of the chromosomal genome (i.e. 29.8-30.2 Mb) was accessible for isolates belonging to the *L. braziliensis* species complex, and 88.1% of the genome (i.e. 28.9 Mb) was accessible for *L. panamensis*. Phylogenetic analyses based on 637,821 single nucleotide polymorphisms (SNPs) revealed a clear distinction between the three *Leishmania Viannia* species and confirmed that *L. peruviana* and *L. braziliensis* correspond to two closely related but distinct monophyletic clades (Fig. 1a). When estimating the proportion of fixed nucleotide differences between species across their combined accessible genome of 28.3 Mb, we found that *L. panamensis* showed fixed differences at 1.06% sites from *L. braziliensis* and 1.21% sites from *L. peruviana*. In contrast, *L. braziliensis* and *L. peruviana* differed at a tenfold lower number of fixed nucleotide differences (0.04%), highlighting their close relatedness.

**Figure 1.**
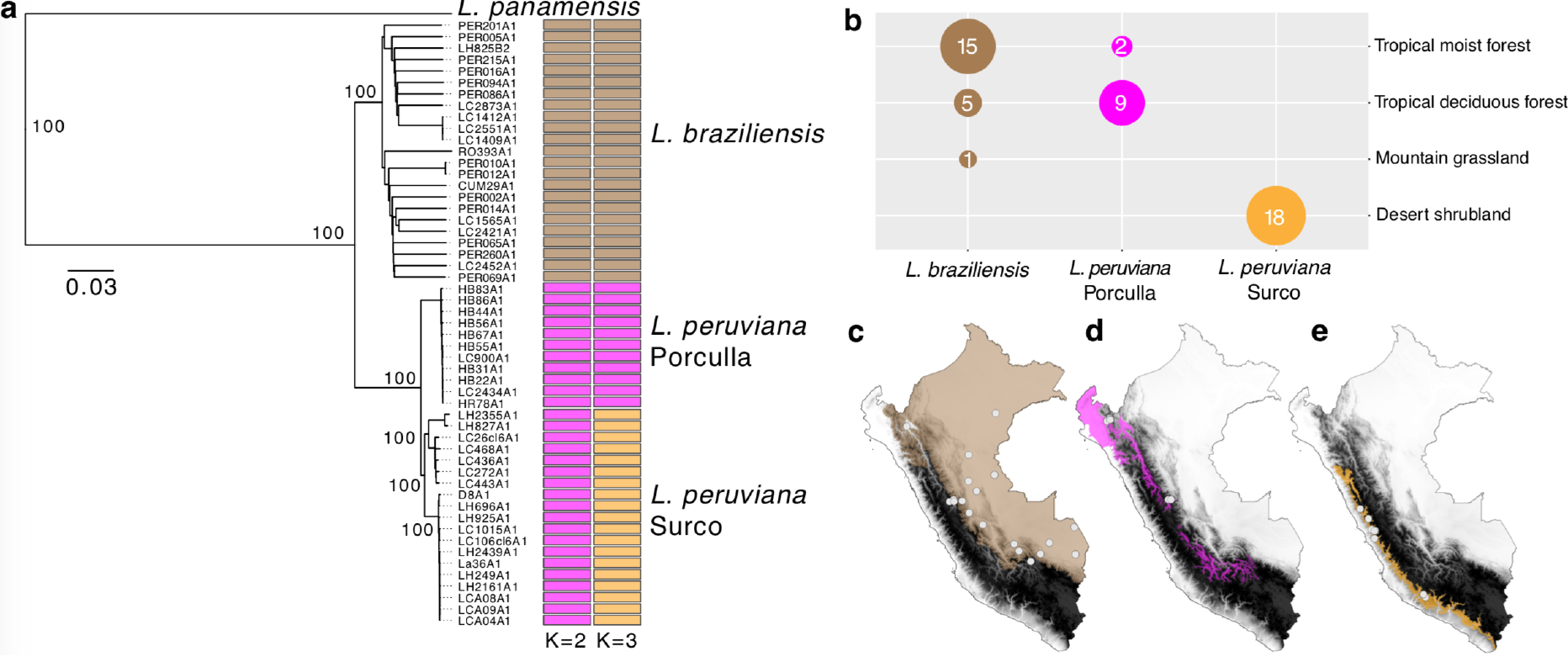
**(a)** Neighbor-Joining phylogenetic tree depicting the genetic ancestry of the *L. braziliensis* species complex (*L. braziliensis* and *L. peruviana*) in Peru including *L. panamensis* as an outgroup. Colored barplots show parasite groups as estimated with ADMIXTURE assuming *K* = 2 and *K* = 3 populations. **(b)** Sample sizes of the three major parasite lineages grouped according their originating biome. **(c,d,e)** Geographic maps of Peru showing the Andean topography in grayscale, the sampling locations of the three major parasite lineages and their corresponding biomes: **(c)** tropical moist forest, **(d)** tropical deciduous forest and **(e**) desert shrubland.

Genotyping across the accessible genome (88.3% - 28.9 Mb) of the 67 *L. braziliensis* species complex isolates disclosed a total of 389,259 SNPs and 114,082 small insertions/deletions (INDELs). SNPs and variable sites (i.e. excluding SNPs fixed in a given species) were evenly distributed across the 35 major chromosomes, but *L. peruviana* showed a 2.6 fold lower density in SNPs (4.1/kb vs. 10.6/kb) and a 9.5 fold lower density in variable sites (1/kb vs. 9.5/kb) compared to *L. braziliensis* (Supp. Fig. 2). While the allele frequency spectrum of *L. braziliensis* was dominated by rare variants, the large majority of SNP loci were entirely fixed (67%) and homozygous (89%) in *L. peruviana* (Supp. Fig. 3), suggesting a strong population bottleneck at the origin of this species. Several of these fixed SNP mutations were virtually absent in *L. braziliensis* and deleterious to genes coding for an ion transporter protein, kinesin-C^30^ and the subunit 2 of the class I transcription factor A complex^31,32^ (Supp. Table 2).

*L. peruviana* and *L. braziliensis* showed a relatively extensive variation in chromosome copy numbers, except chromosome 31 that was tetrasomic for most isolates, and chromosomes 19, 26, 27, 32 and 34 that were disomic for all isolates (Supp. Fig. 4). A high degree of aneuploidy is especially observed among *in vitro* cultivated promastigotes^33^, which may explain the lack of species-specific somy profiles observed here (Supp. Fig. 4, grayscale boxes). A total of 164 gene copy number variations were identified in *L. peruviana* (51 deletions and 113 amplifications). Six deletions and five amplifications were shared among all *L. peruviana* isolates, eight of which encompassed genes encoding cell-surface glycoproteins such as the gp63 leishmanolysin gene family and the ∂-amastin surface glycoproteins (Table 1; Supp. Fig. 5). Other major differences encompass putative proteins encoding kinesin, autophagy protein ATG8 ubiquitin and a glycerol uptake protein (Table 1; Supp. Fig. 5).

**Table 1.** Ancestral deletions and amplifications in montane *L. peruviana* species. The numbers in the two species columns reflect the mean predicted copy numbers with the minimum and maximum copy numbers given between brackets.

| Type | Orthologous Cluster | Product | Chromosomes | *L. peruviana* | *L. braziliensis* |
| --- | --- | --- | --- | --- | --- |
| Amplification | ORTHOMCL29 | delta amastin-like surface protein | 20 | 22 (8-34) | 1 (0-3) |
| Amplification | ORTHOMCL6809 | delta amastin-like protein | bin | 16 (12-21) | 9 (8-12) |
| Amplification | ORTHOMCL75 | Autophagy protein Atg8 ubiquitin like, putative | 19 | 12 (8-16) | 5 (3-7) |
| Amplification | ORTHOMCL66 | kinesin, putative | 20, bin | 18 (15-24) | 12 (10-16) |
| Amplification | - | Amastin | 20 | 13 (10-18) | 8 (6-11) |
| Deletion | ORTHOMCL1 | GP63, leishmanolysin | 10, bin | 37 (23-50) | 64 (53-79) |
| Deletion | ORTHOMCL12 | glycerol uptake protein, putative | 19 | 3 (3-4) | 9 (5-12) |
| Deletion | ORTHOMCL40 | Amastin surface glycoprotein, putative | 20 | 0 (0-1) | 5 (3-7) |
| Deletion | ORTHOMCL6811 | delta amastin-like protein | 8 | 0 (0-1) | 4 (3-5) |
| Deletion | - | Amastin | 20 | 1 (0-2) | 4 (3-6) |
| Deletion | ORTHOMCL6547 | delta amastin | 8 | 0 (0-1) | 3 (1-5) |

### Comparative population genomics of lowland and montane *Leishmania* parasites

Analyses of population structure using unsupervised clustering with ADMIXTURE (Fig. 1a) revealed three major groups of parasites, each corresponding to a particular biome. The first group comprised the lowland *L. braziliensis* parasites that were largely found within tropical moist forests at a median altitude of 631 m (Fig. 1b,c). The second and third group comprised the montane *L. peruviana* parasites found within two different biomes. The Porculla lineage was found at an average altitude of 1,985 m within tropical deciduous forests that span the Huancabamba depression and the North-Eastern slopes of the Peruvian Andes (Fig. 1b,d). The Surco lineage was exclusively found within desert shrubland along the Pacific slopes of the Peruvian Andes, at an average altitude of 2,769 m (Fig. 1b,e). The Surco lineage was further subdivided into three differentiated sublineages, here-after referred to as Surco North (SUN), Surco Central (SUC) and Surco Central/South (SUCS) (Supp. Fig. 6). In contrast, clustering analyses failed to identify subpopulations in *L. braziliensis* (results not shown) and the *L. braziliensis* network featured long branches that separate most isolates with little clade structure, a pattern symptomatic of high gene flow (Supp. Fig. 7).

To make predictions about relative recombination rates, Hardy-Weinberg equilibrium (HWE) was tested by estimating *F*_IS_ per SNP locus in *L. peruviana* and *L. braziliensis* taking into account Wahlund effects (see methods). The *F*_IS_ distribution was skewed towards negative *F*_IS_ for *L. peruviana* (mean *F*_IS_ = −0.54) with almost half of the SNP loci showing *F*_IS_ = −1, suggesting heterozygote genotypes, as would be predicted for a population experiencing predominantly clonal propagation (Supp. Fig. 8a-b; Supp. Table 3). In contrast, *L. braziliensis* displayed a unimodal distribution centered around zero (mean = −0.11), suggesting that the population is close to HWE and that *L. braziliensis* may experience relatively high recombination rates (Supp. Fig. 8c; Supp. Table 3). Despite strong genetic differentiation and reduced recombination rates in *L. peruviana*, there were signals of historical hybridization events, in particular among the Surco populations (Supp. Fig. 9). The occurrence of hybridization among the Surco populations was also highlighted by isolate LH741 that showed a mixed ancestry between the SUN and SUCS populations (Supp. Fig. 6a, 7b).

### Late Pleistocene origin of montane *Leishmania* parasites

The time-resolved phylogeny calibrated based on an assumed substitution rate for maxicircles^34^ suggested that the common ancestor of *L. peruviana* and *L. braziliensis* lived ~128 kya (CI: 85 kya – 175 kya) (Fig. 2a) during the Last Interglacial (130 kya - 115 kya). Subsequent diversification of *L. peruviana* occurred at several occasions during the Last Glacial Period which lasted from 115 kya to 11.7 kya (Fig. 2a). These results suggest that the (sub-)diversification of *L. peruviana* may have been promoted by extensive climatic cycling and vegetational shifts.

**Figure 2.**
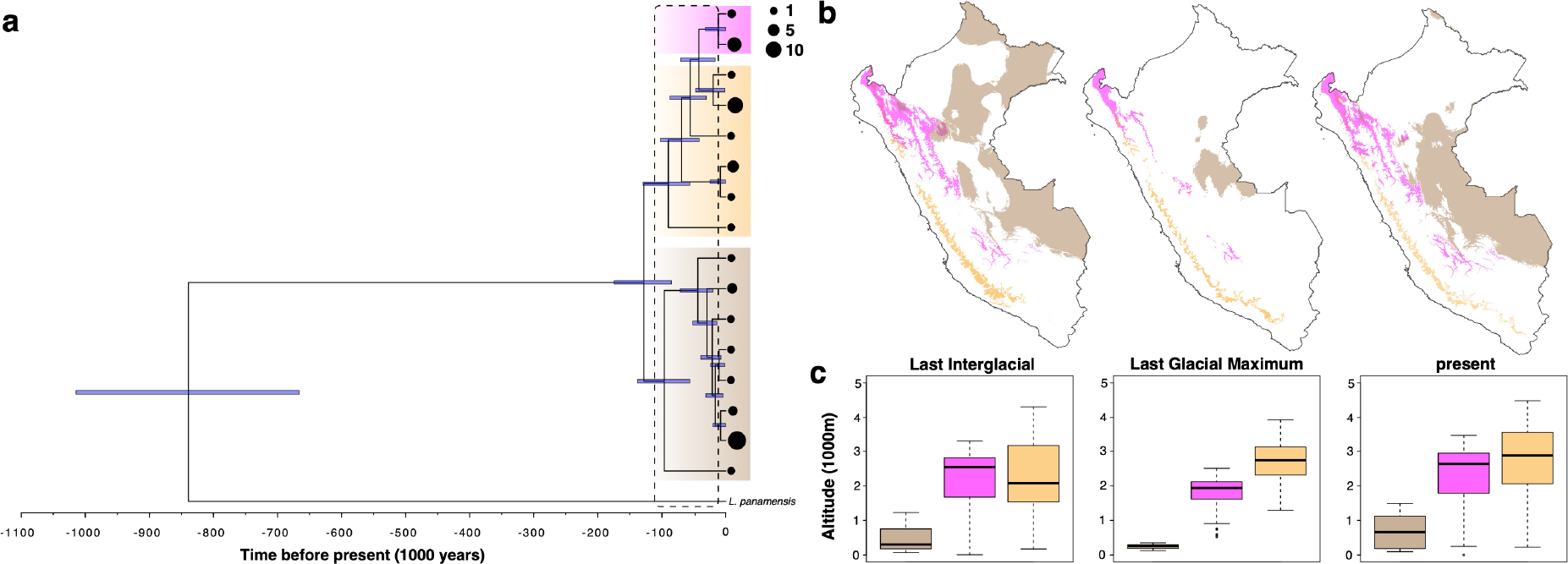
**(a)** Time-calibrated phylogenetic tree for the *L. braziliensis* species complex based on maxicircle gene alignments. Blue node bars represent the 95% highest posterior density of the divergence time estimates. Thick transparent boxes mark the maxicircle haplotypes of *L. braziliensis* (brown), *L. peruviana* Porculla (magenta) and *L. peruviana* Surco (orange). The size of the black circles at the tips of the branches reflect the number of haplotype sequences (legend on the topright). **(b)** Geographic maps of Peru showing the modeled distribution and **(c)** average altitude of the predicted habitat patches, for each of the three major *Leishmania* lineages during the Last Interglacial, Last Glacial Maximum and present (see Fig. 2a for color codes).

Environmental niche modeling (ENM) using present and past bioclimatic variables predicted the putative range of each *Leishmania* population (*L. braziliensis*, *L. peruviana* Porculla and *L. peruviana* Surco) during the Last Glacial Maximum (LGM: 21 kya) and the last inter-glacial (LIG: 130 kya). The bioclimatic variables that contributed most to the model results were temperature and precipitation seasonality. For the present, the predicted areas included the species known distributions in Peru and largely matched the biogeographic regions (tropical moist forest, tropical deciduous forest and desert shrubland) for each *Leishmania* population (Fig. 2b; Supp. Fig. 10). During the LGM, there was a strong contraction, fragmentation and isolation of suitable habitat for all *Leishmania* populations (Fig. 2b; Supp. Fig. 10), resulting in a more pronounced difference in altitude range between *L. braziliensis* and the montane *L. peruviana* lineages (Fig. 2c). During the LIG, the potential range of *L. peruviana* was rather similar, but the range of *L. braziliensis* was slightly shifted to the north of Peru, resulting in a more pronounced spatial overlap with the range of the *L. peruviana* Porculla population (Fig. 2b; Supp. Fig. 10). The average altitude of the habitat patches was considerably different between *L. braziliensis* and the two *L. peruviana* populations.

### Meiotic-like recombination between lowland and montane *Leishmania* parasites

We also included 13 hybrid *L. peruviana* × *L. braziliensis* isolates from the Huánuco region where both *Leishmania* species and their hybrids occur sympatrically^16,17^. Earlier molecular work suggested the presence of four zymodemes and seven microsatellite genotypes within the hybrid population^17^, and here we sequenced isolates from each zymodeme. Of a total 149,735 SNPs that were identified in the hybrid population, 61,804 sites (41.3%) were fixed homozygous and 73,375 sites (49%) were fixed heterozygous, leaving 14,556 segregating sites (9.7%). Heterozygous sites were evenly distributed across the genome and rarely interrupted by homozygous stretches (Supp. Fig. 11). The frequency distribution of allelic read depths at heterozygous sites was centered around 0.5 for all hybrids (data not shown), as would be predicted for diploid *Leishmania* parasites^35^. Hybrids were near-identical, differing by a median 1,245 heterozygous sites and one homozygous site (Supp. Table 4). Exceptions were isolates PER011 and LC2520 that showed 167 homozygous SNP differences (Supp. Table 4), but close inspection revealed that 165 SNPs were located within a 63kb window on chromosome 32 that was homozygous for either parental alleles in the two isolates (Supp. Fig. 11), suggesting that these differences are due to gene conversion events.

Principal Component Analyses (PCA) based on genome-wide SNPs showed that hybrids occupied a tight central position between *L. braziliensis* and the *L. peruviana* SUCS population (Fig. 3a), suggesting that all hybrids are first-generation offspring. Estimates of raw nucleotide differences revealed a similar genetic distance between the hybrids and the *L. braziliensis* isolates LC2551, LC1409 and LC1412 on the one hand, and the *L. peruviana* SUCS population on the other hand (results not shown). In addition, close examination of 89 SNPs identified within the coding region of the mitochondrial maxicircle revealed that all hybrids were identical to *L. braziliensis* isolates LC2551, LC1409 and LC1412 (Fig. 3b). Altogether, these results clearly indicate the following two parent groups of isolates: i) *L. braziliensis* isolates LC2551, LC1409 and LC1412 from the Huánuco region and ii) *L. peruviana* SUCS population from the Lima and Ayacucho regions.

**Figure 3.**
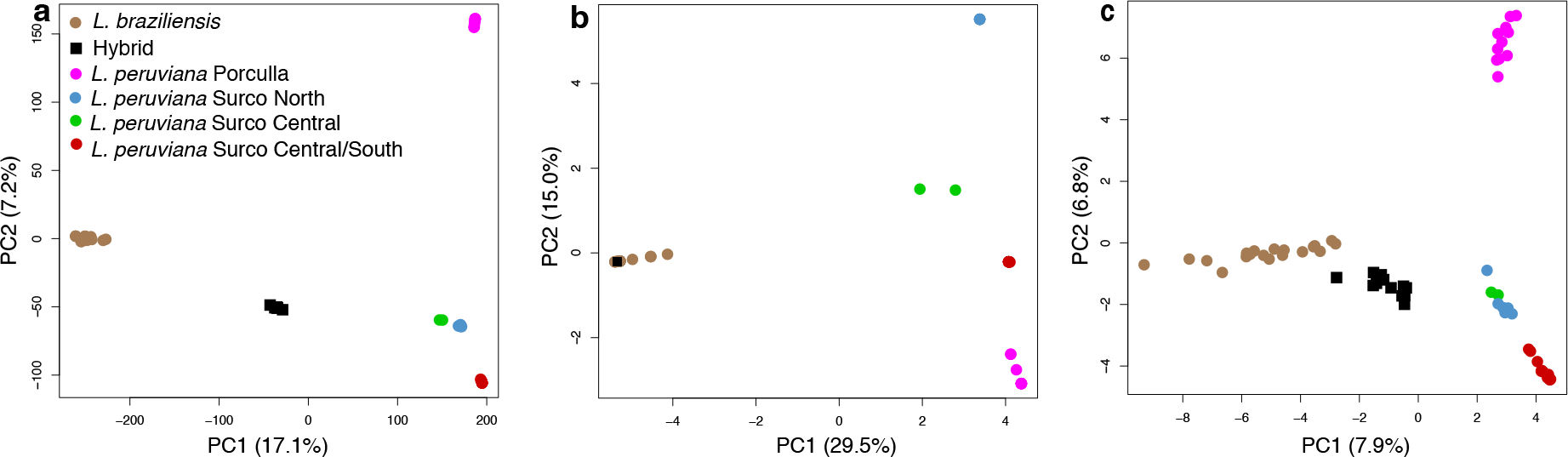
Principal Component Analyses of the *L. braziliensis* species complex in Peru based on 389,259 genome-wide SNPs **(a)**, 89 SNPs of the maxicircle coding region **(b)** and sequence similarity of 950 minicircle sequence classes **(c)**. Hybrid isolates were projected onto the PCA space of *L. braziliensis* and *L. peruviana*, where the first axis separates *L. braziliensis* and *L. peruviana*, and the second axis distinguishes the main *L. peruviana* populations.

To investigate the ancestry of the hybrids in more detail, we focused on the 113,266 SNPs that were shared between the 13 hybrids and representative isolates of each parent group (LH925 for *L. peruviana* SUCS and LC1412 for *L. braziliensis*). SNPs were counted when homozygous for the reference allele (R/R), homozygous for the alternate allele (A/A) or heterozygous (A/R). Of the 54,734 SNPs that were homozygous (A/A) in one parent and absent (R/R) in the other, an average 54,038 SNPs (99.5%) were heterozygous (A/R) in the 13 hybrids. Of the 14,025 SNPs that were homozygous in one parent (A/A) and heterozygous in the other parent (A/R), an average 7,399 SNPs (53%) were heterozygous (A/R) and 6,596 SNPs (47%) homozygous (A/A) in the hybrid offspring. Of the 41,424 SNPs that were heterozygous (A/R) in one parent and absent (R/R) in the other, an average 21,053 SNPs (50.8%) were heterozygous (A/R) and 20,295 SNPs (48.9%) were absent (R/R) in the hybrid offspring. These results confirm that hybrid parasites inherited parental alleles in a 1:1 ratio, as would be predicted for first-generation offspring of a Mendelian cross.

### Biparental inheritance of *Leishmania* mitochondrial minicircle populations

Minicircles were assembled and circularized for each of the 67 isolates using the python package KOMICS (see Methods and Supp. Fig. 12). A total of 9,003 minicircle contigs were assembled for 64 isolates, of which 6,949 (77%) circularized. The assembly process failed for 3 isolates (LC1565, LC1409, LC1412) for which there were insufficient mitochondrial reads. The number of assembled minicircle contigs per isolate did not depend on sequencing depth, as there was no association between median genome-wide read depth and the number of minicircles in *L. braziliensis* (r = 0.20, *p* = 0.41), *L. peruviana* (r = 0.07, *p* = 0.7) and the hybrids (r = 0.03, *p* = 0.92). To validate the quality of the assembly, reads were aligned to the minicircle contigs and several mapping statistics were summarized. First, on average 95% of all mapped reads were properly paired and 93% aligned with a mapping quality larger than 20 (Supp. Table 5). Second, a total of 100 homozygous SNPs were identified within 56 contigs, which is only 0.62% of all contigs, suggesting a robust assembly for the large majority of the minicircle contigs. Third, the length of the majority of the circularized minicircles (6,906 contigs, 99.3%) showed a bimodal distribution around ~740 bp and ~750 bp (Supp. Fig. 13), which is comparable to the minicircle length (~850bp) found in *L. tarentolae*^36^. The remaining 43 contigs (<0.7%) showed twice this length (~1490 bp; data not shown), suggesting that these may be artificial minicircle dimers. Finally, the number of reads containing the Conserved Sequence Block 3 (CSB-3) 12-mer (also called universal minicircle sequence^37^) was calculated as a proxy for the total number of minicircles initially present within the DNA sample. Note that the CSB-3 12-mer is present within both the minicircles^37^ and maxicircles^38^, but here we only used reads that did not align to the maxicircle. On average 95% of all CSB3-containing reads aligned against a given minicircle contig, 90% aligned with a perfect match and 89% aligned in proper pairs (Supp. Table 5), suggesting that KOMICS was able to retrieve the large majority of the minicircles.

Minicircle complexity and ancestry was studied using a clustering approach to find sets of minicircle sequences that show a minimum percent identity with each-other (here-after referred to as minicircle sequence classes or MSCs). These clustering analyses were only done on the circularized minicircle contigs of the expected length, as these would produce the most robust alignments. The number of MSCs decreased sharply from 4,290 MSCs at 100% identity to only 582 MSCs at 95% identity and 311 MSCs at 80% identity (Supp. Fig. 14a). As percent identity decreased, the alignments were more prone to gaps larger than or equal to 2 nucleotides (Supp. Fig. 14b). Specifically, there was a sharp increase in the number of alignments with 3-nt gaps from 97% to 96% identity (Supp. Fig. 14b), suggesting that clustering results may be less robust below the 97% identity threshold. In addition, discriminatory power decreased strongly below the 97% identity threshold as we observed a decrease in the proportion of MSCs unique to *L. peruviana* and an increase in the proportion of MSCs shared between *L. braziliensis*, *L. peruviana* and the hybrids (Fig. 4a). Focusing on the results at 97% identity, a significantly lower number of MSCs were found per isolate in *L. peruviana* (mean = 81 MSCs/isolate) compared to *L. braziliensis* (mean = 147 MSCs/isolate; *p* < 0.0001), with hybrids showing an intermediate value (mean = 111 MSCs/isolate) (Fig. 4b).

**Figure 4.**
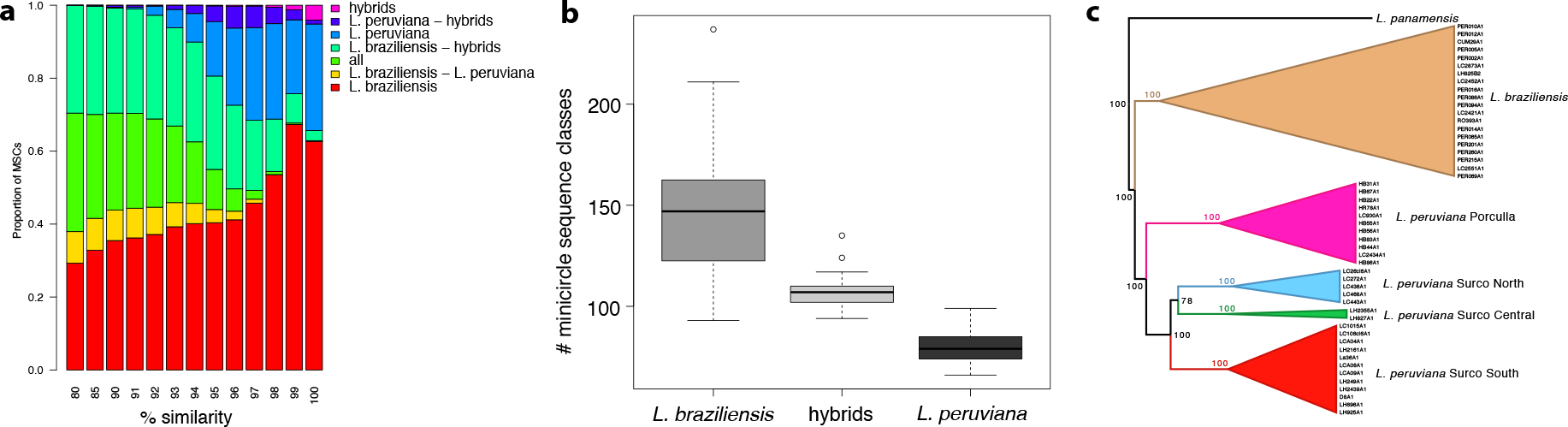
Minicircle complexity and ancestry in the *L. braziliensis* species complex. **(a)** Barplots show the proportion of minicircle sequence classes that are unique or shared between *L. braziliensis*, *L. peruviana* and their hybrids, for each % identity threshold used during the clustering analyses. **(b)** Boxplot showing the number of minicircle sequence classes within *L. peruviana*, *L. braziliensis* and their hybrids **(c)** Neighbor-Joining tree based on a Euclidean distance matrix as estimated based minicircle sequence classes observed in each *Leishmania* isolate.

To study the ancestry of *Leishmania* based on minicircle complexity, we reconstructed a Euclidean distance matrix based on MSCs observed in each *Leishmania* isolate at the 97% identity threshold. A Neighbor-Joining phylogenetic tree including *L. panamensis* revealed a remarkably similar topology as seen using genome-wide SNPs (Fig. 1a), with a clear distinction between the *L. peruviana* Porculla lineage and the *L. peruviana* Surco lineages (Fig. 4c). A PCA based on minicircle absence/presence in each isolate showed an identical pattern as seen with genome-wide SNPs, with the first axis separating *L. braziliensis* from *L. peruviana* and the second axis dividing the main *L. peruviana* populations (Fig. 3c). Interestingly, hybrids did not cluster with either parental species as would be predicted for a uniparentally inherited kinetoplast, but rather occupied an intermediate position between *L. braziliensis* and the *L. peruviana* SUCS population (Fig. 3c). In addition, at 97% identity, virtually all MSCs found in the hybrids are also found within *L. peruviana* and/or *L. braziliensis* (Fig. 4a). When quantifying the number of MSCs shared between each hybrid isolate and either parental species, we found that on average 20.1% of the MSCs in hybrid progeny originated from *L. peruviana* and 69.5% from *L. braziliensis* (Table 2). These results confirm that hybrid parasites contained MSCs unique to each parental species.

**Table 2.** Total number of minicircle sequence classes (MSCs) for the 13 *L. braziliensis* × *L. peruviana* hybrid isolates. Percentages indicate the proportion of MSCs that were unique to *L. braziliensis,* unique to *L. peruviana*, or found in both *L. peruviana* and *L. braziliensis*.

| Isolate | Number of MSCs | *L. braziliensis* | *L. peruviana* | *L. braziliensis* & *L. peruviana* |
| --- | --- | --- | --- | --- |
| LC2877 | 98 | 64.3% | 26.5% | 9.2% |
| HR410 | 111 | 73.0% | 16.2% | 10.8% |
| HR434 | 105 | 76.2% | 14.3% | 9.5% |
| LC2435 | 119 | 73.1% | 16.0% | 10.9% |
| LC2520 | 110 | 74.5% | 15.5% | 10.0% |
| LC1407 | 95 | 64.2% | 24.2% | 11.6% |
| LC1408 | 106 | 65.1% | 26.4% | 8.5% |
| LC1418 | 113 | 65.5% | 24.8% | 9.7% |
| LC1419 | 107 | 66.4% | 22.4% | 11.2% |
| LH1099 | 102 | 72.5% | 16.7% | 9.8% |
| PER011 | 142 | 76.8% | 11.3% | 11.9% |
| LC2851 | 111 | 68.5% | 19.8% | 11.7% |
| HR80 | 124 | 63.7% | 26.6% | 9.7% |
| Average | 111 | 69.5% | 20.1% | 10.4% |

### Mosaic of guide RNA repertoire in hybrid parasites

Putative guide RNA genes (gRNA’s) were identified by aligning minicircle and maxicircle sequences to predicted edited mRNA sequences, allowing for G-U base-pairs^39^ (see Methods). This was done for four isolates representing *L. braziliensis*, *L. peruviana* Porculla, *L. peruviana* SUCS and a hybrid *L. braziliensis* × *L. peruviana* parasite.

All annotated minicircles contained the expected three Conserved Sequence Blocks and 65%-81% (depending on the isolate) had a single predicted gRNA of at least 40 bp complementarity to edited mRNA sequence ~500 bp downstream of the CSB-3 sequence (Supp. Fig. 15, blue dots). Shorter complementary sequences were found throughout the minicircle sequences (Supp. Fig. 15, orange and green dots), suggesting that these were non-specific matches. No putative gRNA genes were identified for 19%-35% of the minicircles (Supp. Table 6). A total of 19-21 gRNAs were identified within the maxicircle of *L. peruviana* and *L. braziliensis* (Supp. Table 7), a number that far exceeded the seven maxicircle-encoded gRNAs (Ma-gRNAs) reported for *L. tarentolae*^36^ and *Crithidia fasciculata*^40^. Five of the Ma-gRNAs identified are shared among all four species, covering editing sites in the 5’-edited maxicircle genes ND7, CYb, A6 and MURF2 (Supp. Table 7; orange bars). The remaining 14-16 Ma-gRNAs identified in *L. braziliensis* and *L. peruviana* were novel and non-redundant candidates, covering editing sites in the pan-edited maxicircle genes ND8, ND9, GR3 and GR4 (Supp. Table 7).

A total of 123 gRNAs were identified in *L. peruviana* Surco, 151 gRNAs in *L. peruviana* Porculla, 154 gRNAs in *L. braziliensis* and 157 gRNAs in hybrid *L. peruviana* × *L. braziliensis* (Supp. Table 6). Paired t-tests showed that there was a significantly lower number of gRNAs between *L. peruviana* Surco on the one hand and *L. peruviana* Porculla (*p* = 0.038), *L. braziliensis* (*p* = 0.049) and the hybrid (*p* = 0.018) on the other hand. The lower number of predicted gRNAs in *L. peruviana* Surco resulted in a lower proportion of editing sites covered by a gRNA (92.52%) when compared to *L. peruviana* Porculla (97.13%), *L braziliensis* (98.17%) and the *L. peruviana* × *L. braziliensis* hybrid (97.37%) (Supp. Table 6).

The distribution and ancestry of the predicted gRNAs was examined across the four pan-edited genes GR3, GR4, ND8 and ND9, revealing two major results (Fig. 5). First, many of the novel Ma-gRNAs covered editing sites that were not covered by minicircle-encoded candidates (Fig. 5, stars), suggesting that these Ma-gRNAs are essential to prevent a break of the 3’-5’ editing cascade. This is most clearly observed for the *L. peruviana* SUCS isolate where Ma-gRNAs covered four different locations in the ND9 gene that were not covered by minicircle-encoded gRNAs (Fig. 5, stars). Second, *L. peruviana* × *L. braziliensis* hybrids showed a mosaic ancestry of gRNAs originating from both parental species, with *L. peruviana* - specific gRNAs aligning in locations where there were no *L. braziliensis* - specific gRNAs (Fig. 5, arrows).

**Figure 5.**
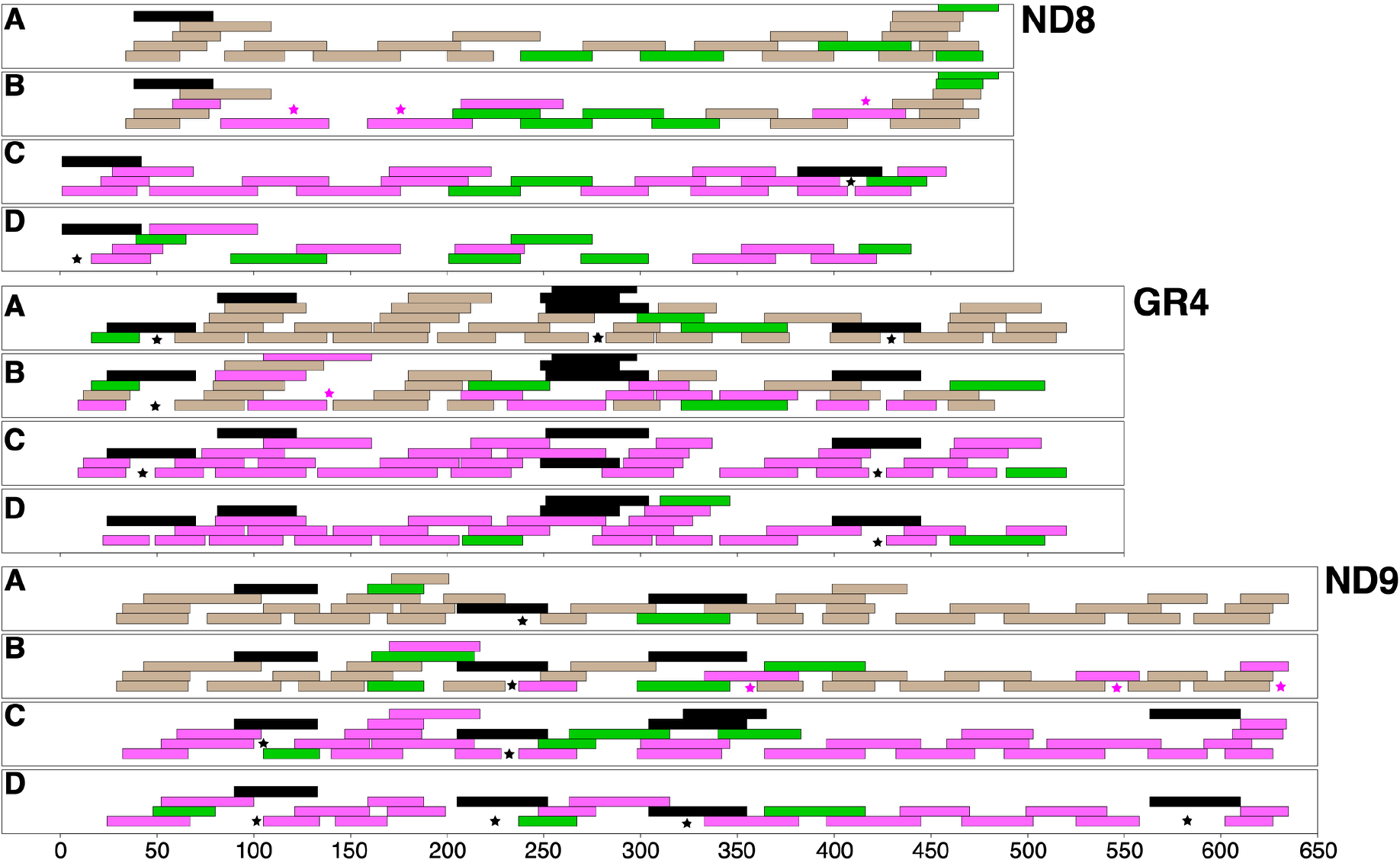
Coverage of editing sites for the pan-edited maxicircle genes ND8, GR4 and ND9. Stacked rectangles are gRNA genes covering editing sites in a given maxicircle gene of *L. braziliensis* isolate LC1412 (A), hybrid isolate HR434 (B), *L. peruviana* Porculla isolate HR78 (C) and *L. peruviana* SUCS isolate (D). Colors indicate whether the gRNA originated from the maxicircle (black) or from a minicircle found in *L. braziliensis* (brown), *L. peruviana* (magenta) or both (green). Black stars indicate editing sites that are covered solely by maxicircle-encoded gRNA candidates. Magenta stars in hybrid isolate HR434 (B) indicate editing sites solely covered by *L. peruviana* gRNA candidates.

## DISCUSSION

We have presented population genomic analyses based on the complete nuclear and mitochondrial genome of natural *Leishmania* isolates that provide a comprehensive view on genetic diversification and subsequent hybridization in parasitic protozoa. Our data supports the major conclusions that 1) ecology is a major driver of reproductive isolation in Neotropical *Leishmania* parasites and 2) meiotic recombination creates mosaic ancestry of mitochondrial genes due to biparental inheritance of minicircles.

Andean *L. peruviana* parasites demonstrated stable long-term genetic diversification, evolving as near-clonal lineages that emerged from admixed Amazonian ancestors. The origin of the Andean lineages was accompanied by a strong population bottleneck, as evidenced by a genome-wide fixation of SNP polymorphisms. Several of these fixed SNP mutations were deleterious to genes involved in the parasite’s biosynthetic capability or cytokinesis, which may explain the slower growth rate *in vitro* of L. *peruviana* compared to *L. braziliensis*^41,42^. In addition, all *L. peruviana* isolates shared eight structural variations encompassing genes involved in host-parasite interactions. Most notably was a 51.1 kb deletion spanning the gp63 leishmanolysin gene family, important for amastigotes to subvert the macrophage immune response^43^. Reduced copy numbers may explain the lower virulence of *L. peruviana* amastigotes *in vivo*^42^.

Beside a clear dichotomy in population genomic structure between the Andean and Amazonian *Leishmania* species, parasite genomic diversity was principally partitioned by ecotype. The strict ecological association of the major *Leishmania* lineages in Peru suggest that reproductive isolation occurred through ecological fitting, a process whereby an organism colonizes and persists in a new or modified environment^44^. Such exaptation into novel environmental niches is often accompanied by host switches and facilitated by climatological variation and ecological perturbation^45^. Here we show that the history of diversification of Andean lineages is limited to the Late Pleistocene and intimately associated with vegetation changes. We propose that spatial overlap of Andean and Amazonian ecotypes during the Last Interglacial may have facilitated the intrusion of Amazonian *Leishmania* parasites into the Andean ecotypes containing different sandfly vector communities^46,47^, with subsequent habitat and altitudinal range contractions during last glacial episodes fueling the diversification of the major Andean lineages. Together with observations in trypanosomes^48,49^, our data suggests a key role for ecological fitting in delineating major trypanosomatid parasite clades.

In the Eastern Andean valley of the Huánuco region, signs of meiotic recombination between the two *Leishmania* species are markedly clear. All *L. peruviana* × *L. braziliensis* hybrids contained both parental alleles, providing evidence that hybrids inherited a full set of chromosomes from each parent and are thus full genome hybrids. The observation of near-identical hybrid genomes sampled over a period of 11 years suggest that hybrids arose from a rare mating event between *L. peruviana* and *L. braziliensis*, although we cannot exclude the possibility of multiple mating events including closely related parental parasites. The high number of heterozygous sites across the genome and the central position of the hybrids between either parental species in the PCA space provide clear evidence that they are natural first-generation hybrids that propagated mitotically since the initial cross. The absence of genomic signatures of backcrossing^24,50^ or inbreeding^26^ suggest that these *L. peruviana* × *L. braziliensis* hybrids are sterile, as shown experimentally for inter-species *Leishmania* hybrids^24^. Hybrid sterility among closely related species is the most common form of postzygotic reproductive isolation^51^–^53^, and may be one of the factors that contributed to the speciation of *L. peruviana*.

The mitochondrial minicircles were inherited from both parental species, a phenomenon that has only been described sporadically in Trypanosomatids^54–56^. In contrast, the mitochondrial maxicircles demonstrated clear and consistent uniparental inheritance from the *L. braziliensis* parent. While these observations would suggest differential inheritance of maxi- and minicircles, the most likely explanation is that the kinetoplast DNA networks fused during genetic exchange into a single hybrid network that homogenized during subsequent mitotic divisions. Parallel observations in trypanosomes support this hypothesis. In *T. brucei*, experimental hybrid progeny contained minicircle^54,55^ and maxicircle^57,58^ types of the two parents, whereby maxicircles rapidly homogenized to either parental types during subsequent mitotic divisions^58^. Assuming a stochastic segregation model, it was estimated that fixation in experimental *T. brucei* clones would complete in 139 generations (~35 days) for 50 maxicircles and 27,726 generations (~19 years) for 10,000 minicircles^58^. In *Saccharomyces cerevisiae*, the mitochondrial genome homogenizes in only 20 generations^59^. Rapid stochastic loss of maxicircles following mating could explain why there is currently no evidence for maxicircle heteroplasmy in natural or experimental *Leishmania* hybrids. Nevertheless, the presence of minicircles from both parental species increased the complexity of mitochondrial genomes in hybrid parasites. We show that predicted guide RNA genes originating from both *L. peruviana* and *L. braziliensis* aligned to the *L. braziliensis* maxicircle, resulting in mosaic maxicircle genes. Hence, biparental inheritance of minicircles may present parasites the opportunity to incorporate novel combinations of guide RNA genes, some of which might provide more efficient editing than others, potentially increasing parasite fitness.

Our observations significantly expand our appreciation on the genetic consequences of hybridization in parasitic protozoa, and insinuate that recombination may be crucial to their long-term survival in the wild^36^. Mathematical models in *L. tarentolae* showed that non-essential minicircle classes were lost within a few hundred generations in the absence of recombination^60^. This loss may be especially efficient in *Leishmania* species that contain a single gRNA per minicircle class. Here, the number of minicircle classes was significantly lower within the near-clonal and bottlenecked Andean parasites when compared to the admixed Amazonian *Leishmania* populations, suggesting that minicircle complexity is shaped by parasite population biology. Within this context, an interesting observation is the concordant population structure as observed with genome-wide SNPs and mitochondrial minicircles, while the mitochondrial maxicircle revealed different ancestries. This observation clearly suggests that minicircles follow a similar evolutionary path as the nuclear genome, which may indicate that compatibility between nuclear genes and minicircle-encoded guide RNA genes is essential to maintain efficient respiration. Hence, beside replenishing the minicircle repertoire, biparental inheritance of minicircles may allow for selection in recovering optimal mito-nuclear interactions.

## METHODS

### Parasite collection and DNA sequencing

We included a total of 31 *L. peruviana* isolates from Peru, originating from the regions of Piura (N=9), Huánuco (N=2), Ancash (N=6), Lima (N=6) and Ayacucho (N=6) that largely reflect the distribution of Andean CL (Supp. Fig. 1; Supp. Table 1), with one Peruvian isolate of unknown origin. For comparative purposes, we also included 23 *L. braziliensis* isolates from Peru (N=21), Bolivia (N=1) and Brazil (N=1). In Peru, *L. braziliensis* isolates originated from the regions of Huánuco (N=8), Cajamarca (N=1), Cusco (N=3), Madre de Dios (N=3), Loreto (N=2), Junín (N=1), Pasco (N=1) and Ucayali (N=2) (Supp. Fig. 1; Supp. Table 1). We also included 13 isolates previously characterized as hybrid *L. braziliensis* × *L. peruviana* (Dujardin et al. 1995; Nolder et al. 2007) from the Huánuco region where both *Leishmania* species occur sympatrically. For comparative purposes, we also included *L. panamensis* isolate REST417. Parasites were grown in liquid culture medium for 3-4 days at the Antwerp Institute of Tropical Medicine or the London School of Hygiene and Tropical Medicine, and their DNA was extracted using either the QIAGEN QIAamp DNA Mini Kit or a phenol-chloroform extraction. Paired-end sequencing (2×100bp or 2×150bp) was performed on Illumina HiSeq at the Wellcome Sanger Institute.

### Mapping and variant calling

Paired sequence data were aligned against a novel long-read assembly of the M2904 reference genome containing the 35 major chromosomes that cover 32.73Mb and a complete circularized maxicircle sequence of 27.69kb. Mapping was done using SMALT v0.7.4 (https://www.sanger.ac.uk/science/tools/smalt-0), whereby the hash index was built with words of 13 base pair length (k=13) that are sampled every other position in the genome (s=2). Duplicate reads were tagged using MarkDuplicates as implemented in Picard tools v1.92 (https://broadinstitute.github.io/picard/).

SNP and small INDEL calling was done in GATK v4.0.2^61^. More specifically, we used GATKs HaplotypeCaller to produce genotype VCF files for every isolate, CombineGVCFs to merge the genotype VCF files of all isolates, GenotypeGVCFs to perform joint genotyping and finally SelectVariants to separate SNPs and INDELs. Low-quality SNPs were excluded using VariantFiltration when QUAL < 500, DP < 5, QD < 2.0, FS > 60.0, MQ < 40.0, MQRankSum < −12.5 or ReadPosRankSum < −8.0, or when SNPs occurred within SNP clusters (clusterSize = 3 and clusterWindowSize = 10). Low-quality INDELs were excluded when QD < 2.0, FS > 200.0 or ReadPosRankSum < −20.0. Our SNP and INDEL datasets were further refined by running GATK CallableLoci on each sample BAM file to determine genomic intervals that are callable in each sample with the following parameters: --minDepth 5 –minBaseQuality 25 –minMappingQuality 25. BEDOPS^62^–intersect was used to identify the callable genomic regions common to all isolates, and only variants within the callable genome were retained for downstream analyses. The final set of SNPs and INDELs were annotated using the *L. braziliensis* M2904 annotation file with SNPEFF v4.3^63^.

### Population genomic and phylogenomic analyses

Population genomic structure was examined using phylogenetic network analyses in SPLITSTREE v4^64^, Principle Component Analyses in ADEGENET^65^ as implemented in R v3.5.1^66^, and genotype-based clustering analyses in ADMIXTURE v1.3.0^67^. Weir and Cockerham’s *F*_ST_ were estimated in non-overlapping 50kb windows using VCFtools v0.1.12^68^. Neighbor-Joining trees were build with the R package APE^69^ based on raw nucleotide distances (for genome-wide SNPs) or Euclidean distances (for minicircles; see below).

To test for Hardy-Weinberg Equilibrium (HWE), *F*_IS_ was calculated in R using the formula 1-(Ho/Hs) where Ho is the observed proportion of heterozygous genotypes and Hs the expected proportion of heterozygous genotypes assuming HWE (i.e. 2*pq* with *p* and *q* the frequency of the reference and alternate alleles, respectively). To control as much as possible for any spatio-temporal Wahlund effects, analyses of HWE were focused on four *L. peruviana* isolates sampled in 1989-1990 in the Piura region, five *L. peruviana* isolates sampled in 1990 in the Ayacucho region, and six *L. braziliensis* isolates sampled between 1991-1995 in the Huánuco region.

Mitochondrial phylogenetic analyses were specifically focused on five maxicircle genes that are never edited (COI, ND1, ND2, ND4 and ND5) in order to guarantee robust alignments devoid of INDELs, and with clear start and end positions. GATK’s FastaAlternateReferenceMaker was used to incorporate strain-specific SNPs into the mitochondrial gene sequences of each isolate. Note that we could not find any heterozygous SNP, suggesting absence of heteroplasmy, and indicating that all *L. peruviana* isolates contained a single maxicircle sequence type. Gene alignments were concatenated and identical sequences (i.e. haplotypes) were removed using the R package APE, resulting in a final dataset of 17 unique haplotypes of length 7,005bp. A dated phylogeny was obtained in BEAST v1.8.2^70^ with a strict molecular clock, a HKY substitution model and a Yule tree prior. For molecular clock analyses, we could not resort to tip dating as mitochondrial mutation rates are too slow to generate sufficient mutations in our sampling time range. We therefore assumed a strict molecular clock of 0.8% per Myr^34^.

### Ecological Niche Modeling

Species distribution models of the three studied taxa were generated using the program MAXENT v.3.3.3^71^, which estimates the potential distribution based on presence-only data. MAXENT is particularly well suited for species with few data records^72^. Environmental layers consisted of 19 temperature and precipitation variables downloaded from the WorldClim data set (http://www.worldclim.org) and past climate reconstructions at a scale of 30 arc-seconds (ca. 1km^2^) for current and Last interglacial (120-140kya) scenarios and 2.5 arc-minutes (c. 5km^2^) for Last Glacial Maximum (21kya).

Current climatic data from each occurrence point were extracted using the program Quantum-GIS v2.18 (http://www.qgis.org). For *L. braziliensis*, we only included samples from the tropical moist forest below 700m. Highly correlated variables (*R* > 0.7) were removed to avoid overfitting the data. The 19 bioclimatic variables were reduced to six variables for *L. braziliensis*, two variables for *L. peruviana* Porculla and six variables for *L. peruviana* Surco. The models were run with the following parameters: quadratic, product, threshold and hinge, 500 iterations, regularization multiplier equal 1 and 10 replicates subsampled. We used the average prediction from all the model replicates to construct the ENM species distribution maps. Present-day ENMs were projected into bioclimatic variables predicted for two different past scenarios, Last Interglacial (LIG; 120–140 kya) and LGM (21 kya) from MIROC (Model of Interdisciplinary Research on Climate) and CCSM3 (Community Climate System Model) models. A jackknife procedure was performed to measure the percentage of contribution and the importance of the variables to the models.

### Chromosome and gene copy number variation

Chromosome and local copy number variations were calculated based on haploid read depths. Per site coverages were obtained with SAMTOOLS v1.0^73^. Assuming diploidy, aneuploidy was estimated as two times the haploid chromosomal read depths, which was obtained by normalizing the median chromosomal read depths by the median genome-wide read depth. *L. peruviana-*specific local copy number variations were detected by comparing haploid read depths in non-overlapping 10kb windows or across protein-coding genes between *L. peruviana* and *L. braziliensis*. Haploid copy numbers were obtained by normalizing the median read depths per 10kb window or per gene by the median chromosomal read depth. The haploid copy numbers of genes were summed up per orthologous gene group. In order to identify structural variations specific to *L. peruviana*, we subtracted the haploid copy numbers per *L. peruviana* isolate by the average haploid copy number observed in *L. braziliensis*, yielding a normal distribution centered around zero for each *L. peruviana* isolate. Local copy number variations were then defined where the z-score was lower than −3 (deletions) or larger than 3 (amplifications). The haploid copy numbers of consecutive deletions/amplifications in 10kb windows were summed up to obtain an estimate of the total haploid copy number within a given genomic region.

### Automated assembly and circularization of mitochondrial minicircles

We implemented a novel bioinformatics pipeline in python to automate the assembly and circularization of minicircle sequences from short-read whole genome sequence data. An overview of the pipeline named KOMICS (Kinetoplast genOMICS) is given in Supp. Figure 12, and the program is available at https://github.com/FreBio/komics.

KOMICS takes as input paired-end reads in BAM format, whereby reads were aligned to a nuclear reference genome. Reads that did not align to the nuclear genome are extracted from the alignment file. In order to minimize bias in the assembly process due to sequencing errors, reads are quality trimmed using TRIMMOMATIC v0.32^74^ with the following settings: bases with a quality below 30 are cut from both ends of the read (LEADING:30 TRAILING:30), reads are cut once the average quality within a 10bp window drops below 30 (SLIDINGWINDOW:10:30), reads below 100bp in length are dropped (MINLEN:100) and Illumina adapters were removed (ILLUMINACLIP:TruSeq3-PE.fa:2:30:15:1). High quality reads are then used for *de novo* assembly using a multiple *k-*mer strategy in MEGAHIT^75^, currently the most efficient assembler optimized for large and complex metagenomics sequencing data. As MEGAHIT deals with non-uniform sequencing depths^75^, it is suitable for assembling minicircle sequences that show a large variability in copy numbers.

The resulting contigs of the MEGAHIT assembly are filtered for the presence of the Conserved Sequence Block 3 (CSB3), a 12-bp minicircle motif, also known as the universal minicircle sequence, that is highly conserved across all Kinetoplastida species^37^. By default, KOMICS uses the known CSB-3 motif GGGGTTGGTGTA and its reverse complement to extract contigs of putative minicircle origin. For circularization, komics uses BLAST^76^ as a strategy to identify a sequence that is in common at the start and the end of a given minicircle contig. MEGABLAST is run on the entire set of minicircle contigs with the low complexity filter turned off and allowing a maximum e-value of 10^−5^. The BLAST output is processed to retain only hits among the same minicircle contig (avoiding artificial dimers) with 100% identity and a minimum 20bp overlap at the start and end of a given contig. Whenever an overlap is found, the contig is classified as circular and the duplicated sequence at the start of the contig is removed. Finally, the assembly is polished by putting the CSB3-mer at the start.

The quality of the minicircle assembly is verified by aligning the unmapped reads to all minicircles using SMALT, whereby the circular minicircles were extended by 150bp, and estimating the following mapping metrics. First, the quality of the total assembly for a given sample is verified by estimating the number of reads that are (i) mapped, (ii) perfectly matched, (iii) properly paired (including reads that align at both ends in opposite direction) or (iv) aligned with a minimum mapping quality of 20. The same numbers are also estimated but only for those reads that contain the CSB3-mer, allowing to estimate the proportion of successfully assembled minicircles for a given sample. Second, the quality of each minicircle assembly is verified by estimating the same metrics as listed above for the total assembly, and by estimating the mean, median, minimum and maximum read depth.

### Prediction of guide RNA genes on mitochondrial maxicircles and minicircles

Genome and transcriptome sequencing data were generated for *L. braziliensis* LC1412, *L. peruviana* HR78 and hybrid *L. braziliensis* × *L. peruviana* isolate HR434. Genomic reads were aligned to the *L. braziliensis* reference genome using SMALT as described above, but this time with the maxicircle masked. Unmapped reads were extracted and used for assembling the mitochondrial maxicircles and minicircles. Putative maxicircle contigs were generated with SPADES v3.13^77^ using a multiple *k*mer strategy (21, 33, 55, 77, 99, 127), identified by BLAST and annotated with RATT^78^ using the *L. tarentolae* annotation file. This resulted in a 18,775bp maxicircle fragment for *L. peruviana*, 18,314bp for *L. peruviana* LCA04 and 19,988bp for *L. braziliensis* LC1412, all covering the maxicircle coding region. Minicircles were assembled with KOMICS.

In order to obtain species-specific differences in edited maxicircle mRNA sequences, A, C and G residues in published sequences of *L. tarentolae*^79–82^ and/or *L. mexicana amazonensis* LV78^83^ were corrected based on the assembled maxicircle sequences. Illumina reads from whole-cell RNA sequencing were then aligned to these manually edited sequences using SMALT, and the alignments were carefully inspected for indications of potential differences from the published editing patterns. Canonical gRNAs were then predicted as follows. Coding and template strands of maxicircles and template strands of minicircles were aligned (<u>not</u> permitting gaps) to edited mRNA to predict canonical gRNAs. Each strand was split into 120-nt fragments with each fragment overlapping by 60 nt. Given that gRNA genes are about 40 nt long, an overlap of 60 nt was deemed sufficient to capture all gRNAs. Each fragment was then aligned to all edited mRNA sequences. All unique alignments which met the following criteria were recorded:

- contained from 25 (for minicircles) and 40 (for maxicircles) to 60 non-contiguous matches (Watson-Crick or G-U basepairs)
- contained no more than two contiguous mismatches
- had an anchour duplex of at least 6WC basepairs or 5WC+1GU+1WC bps or 4WC+1GU+2WC bps
- covered at least one U insertion or deletion event.

Plots of predicted gRNA positions on minicircles (Supp. Fig. 15) revealed that highly likely gRNAs (i.e., those longer than 40nt with low frequency of mismatches) occurred at a well-defined position of between 450 to 525 nt downstream from the start of CSB3. All gRNAs falling into this region were assumed to be canonical gRNAs. This does mean that some minicircles encode more than one predicted canonical gRNA. In such cases, without transcriptomic data, it is impossible to determine which predicted gRNAs are transcribed.

