## Supplementary Figures and Tables for "Ecological divergence and hybridization of Neotropical *Leishmania* parasites"

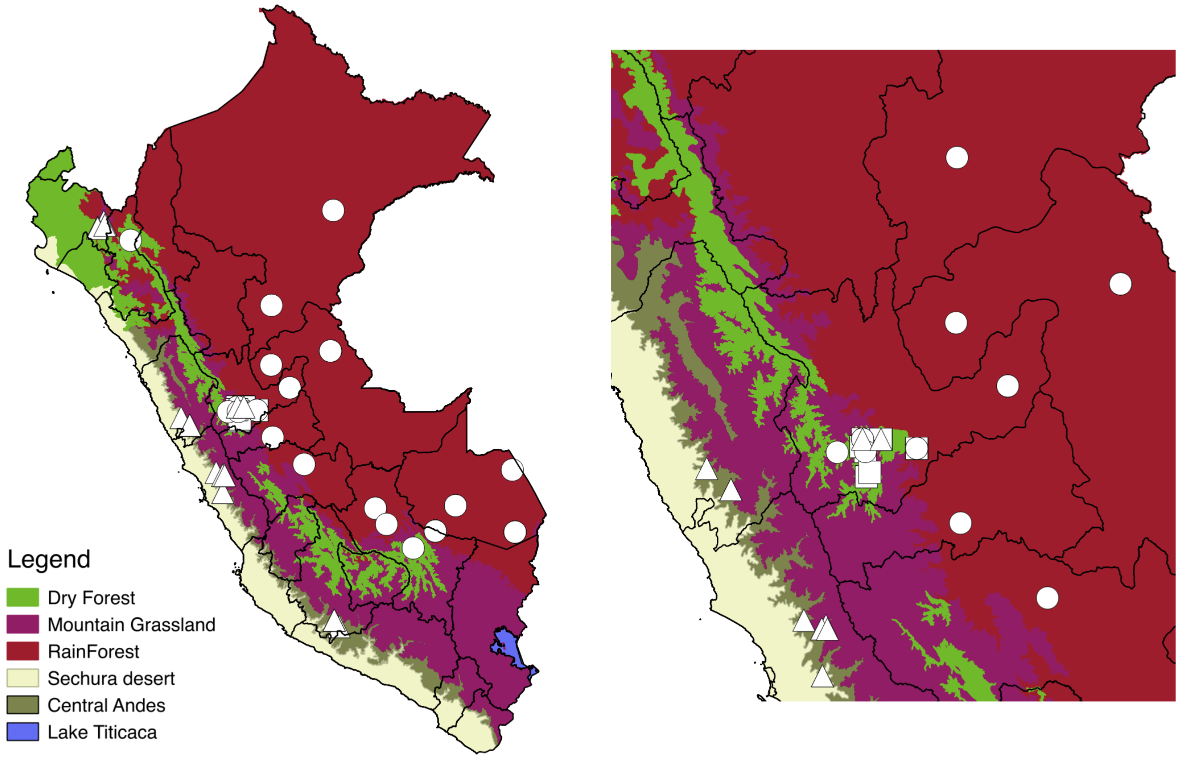

**Supplementary Figure 1.** Map of Peru depicting the main biogeographic regions (see legend), the administrative regions (black lines) and the distribution of isolates of *L. peruviana* (triangles), *L. braziliensis* (circles) and their hybrids (squares). Note that symbols overlap and the size of the symbols does not reflect the number of isolates sampled at a given geographical location. Map on the right focuses on the Huánuco region where *L. braziliensis, L. peruviana* and their hybrids occur sympatrically.

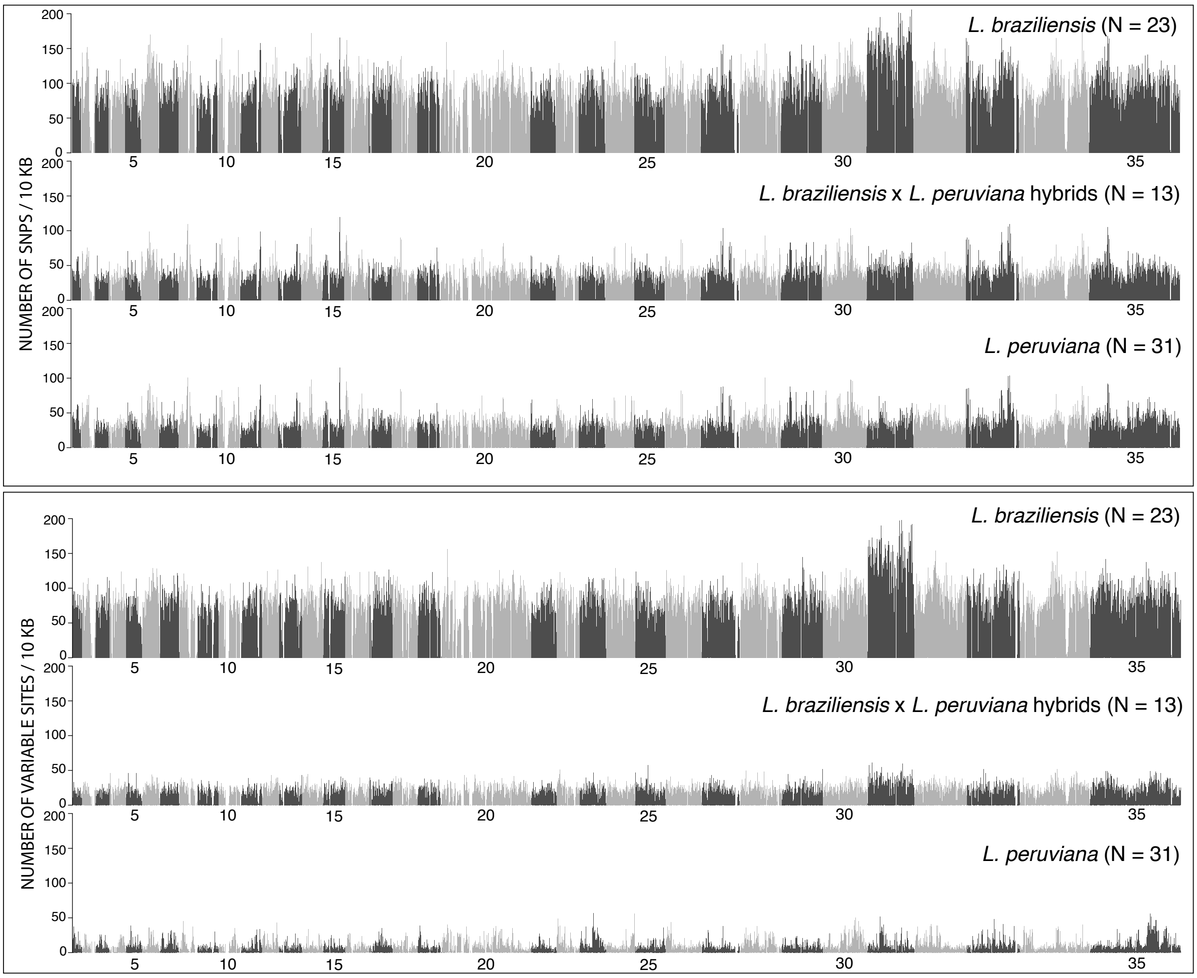

**Supplementary Figure 2.** Number of SNPs (top panel) and variable sites (bottom panel) were estimated within 10kb genomic windows for 23 *L. braziliensis*, 31 *L. peruviana* and 13 hybrid isolates. Genome-wide median numbers are shown on the right of each barplot. Numbers below barplots reflect chromosome numbers.

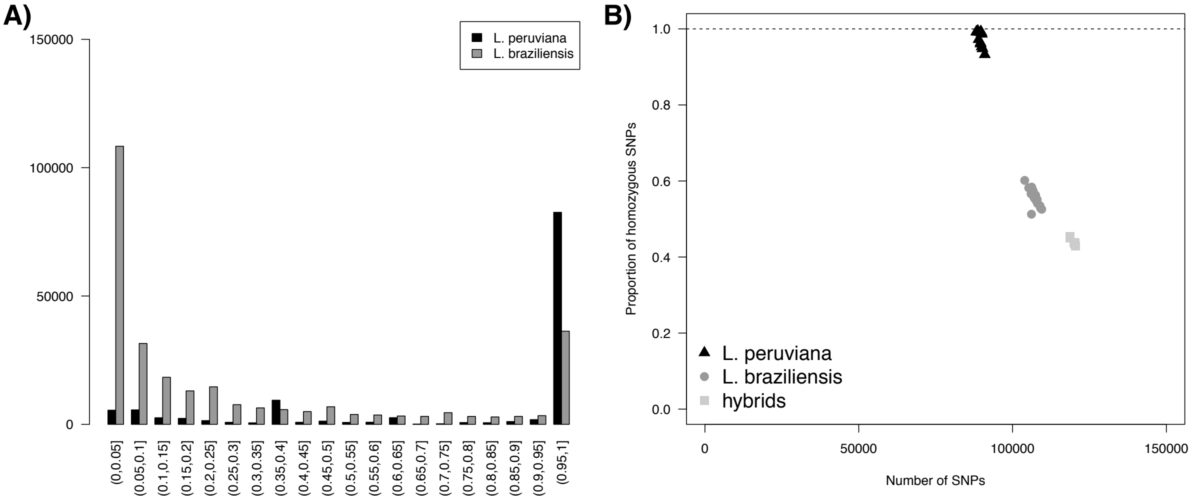

**Supplementary Figure 3. (a)** Alternate (non-reference) allele frequency distribution for *L. braziliensis* and *L. peruviana.* **(b)** Proportion of homozygous SNPs versus the number of SNPs per *L. peruviana*, *L. braziliensis* or hybrid genome.

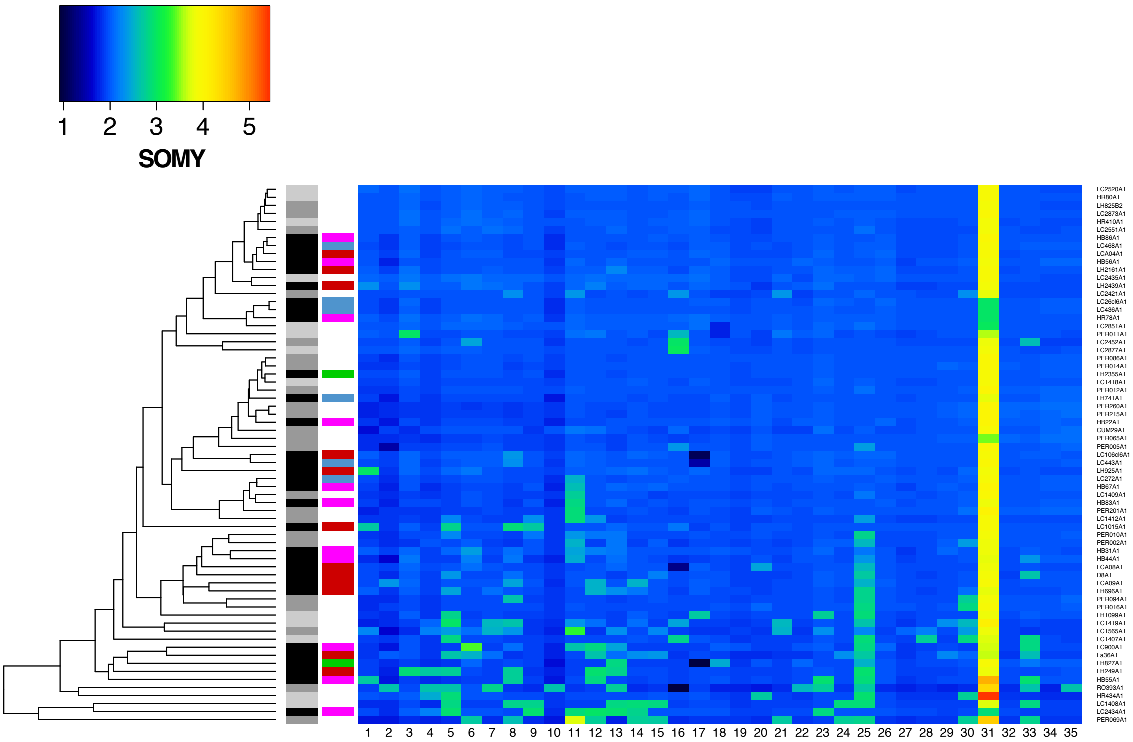

**Supplementary Figure 4.** Somy variability within the *L. braziliensis* species complex for each of the 35 major chromosomes. Dendrogram on the left clusters parasites with similar somy estimates, with aneuploid parasites appearing at the bottom of the heatmap and diploid parasites at the top of the heatmap (with exception of the tetrasomic chromosome 31). Grayscale boxes at the tips of the dendrogram represent *L. peruviana* (black), *L. braziliensis* (dark grey) and their hybrids (light grey). The coloured boxes at the tips of the dendrogram represent the different *L. peruviana* subpopulations (see Supp. Figure 6).

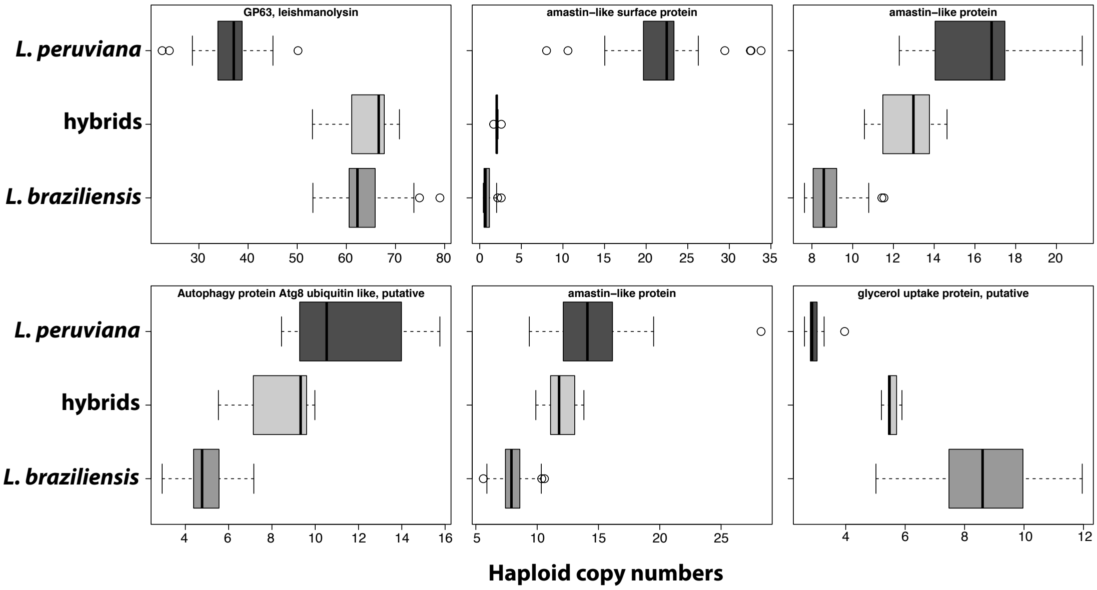

**Supplementary Figure 5. Major structural variations in the *Leishmania braziliensis* species complex.** Boxplots summarize the haploid copy numbers of orthologous gene groups that were found to be most significantly different between *L. peruviana* and *L. braziliensis*.

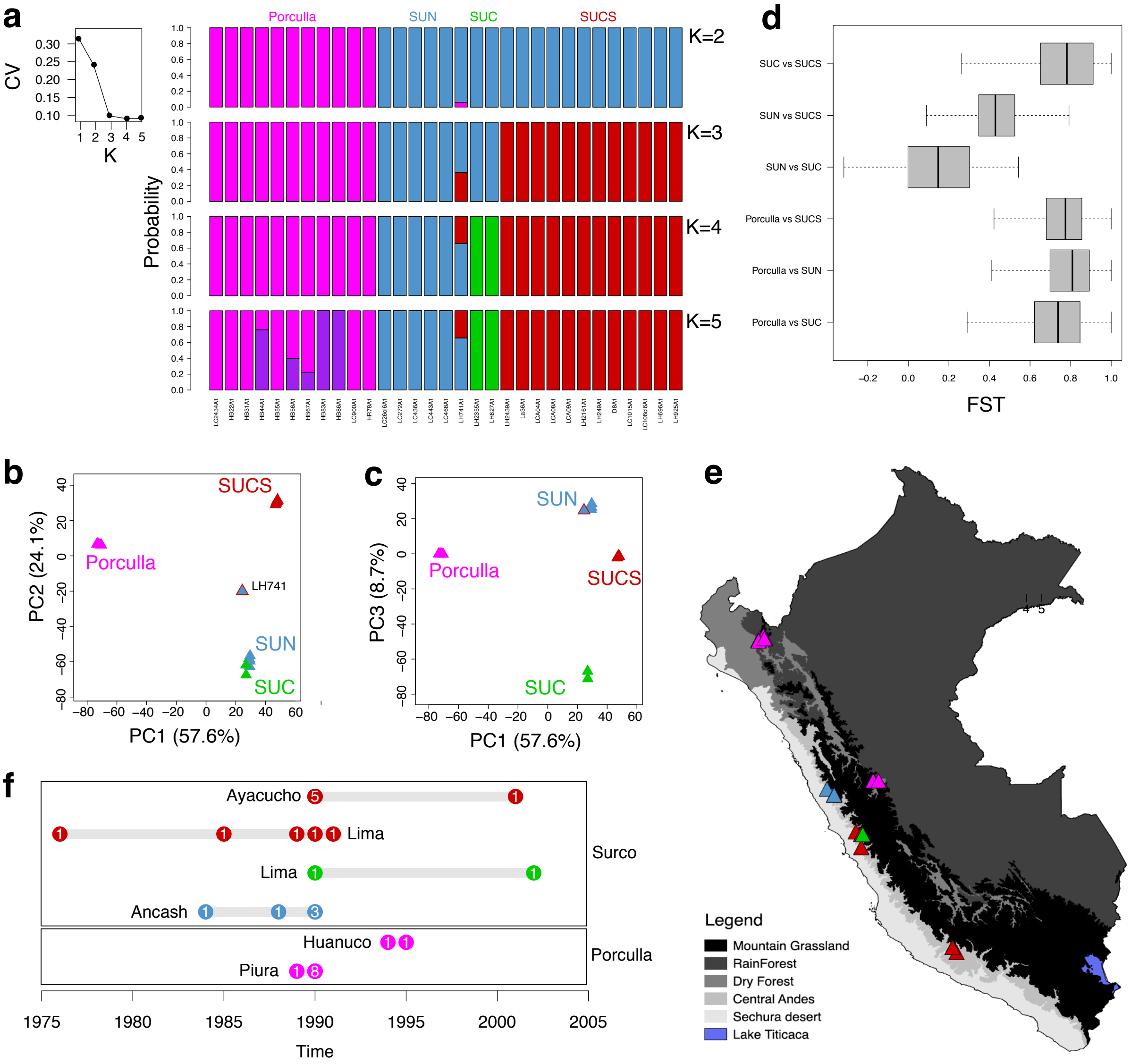

**Supplementary Figure 6. (a)** Analyses of population structure in ADMIXTURE for *L. peruviana* based on 148,246 genome-wide bi-allelic SNPs. Line plot shows the low cross-validation error (CV) for $K = 1$ to $K = 5$. Barplots summarize the membership probabilities for $K = 2$ to $K = 5$. **(b-c)** Principle Component Analyses reveals four groups of parasites (Porculla, SUN, SUC and SUCS) that are separated by the first three principle components explaining a total 90.4% of the variability. Isolate LH741 is a putative hybrid and occupied a central position between the SUN/SUC and the SUCS population. **(d)** Boxplot summarizing genome-wide $F_{ST}$ estimates as calculated within discrete windows of 50kb between the *L. peruviana* populations. **(e)** Geographic map of Peru depicting the main ecogeographic regions and the locations of all *L. peruviana* isolates. **(f)** Sample sizes for *L. peruviana* according to province, population and time, revealing spatio-temporal stability of population structure in *L. peruviana*.

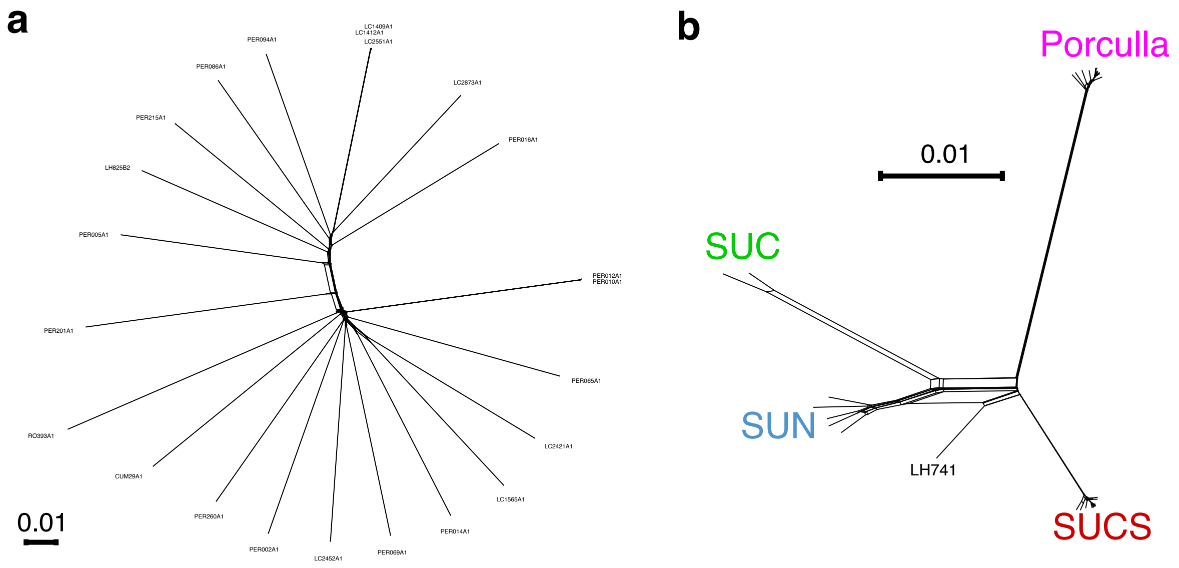

**Supplementary Figure 7.** Phylogenetic network analyses based on uncorrected *p*-distances for *L. braziliensis* **(a)** and *L. peruviana* **(b)**.

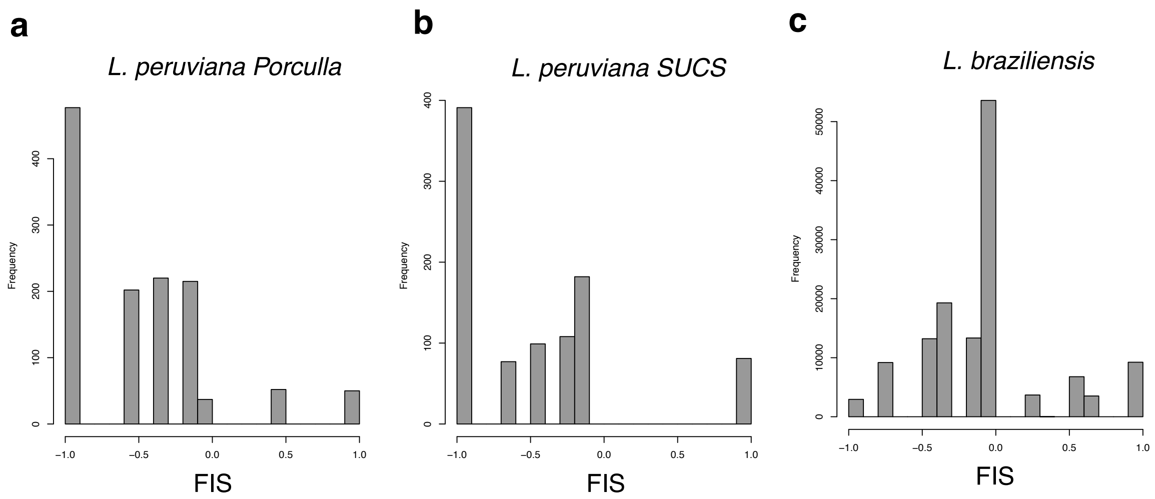

**Supplementary Figure 8.** Distribution of $F_{IS}$ as estimated per locus for each of the three major *Leishmania* populations.

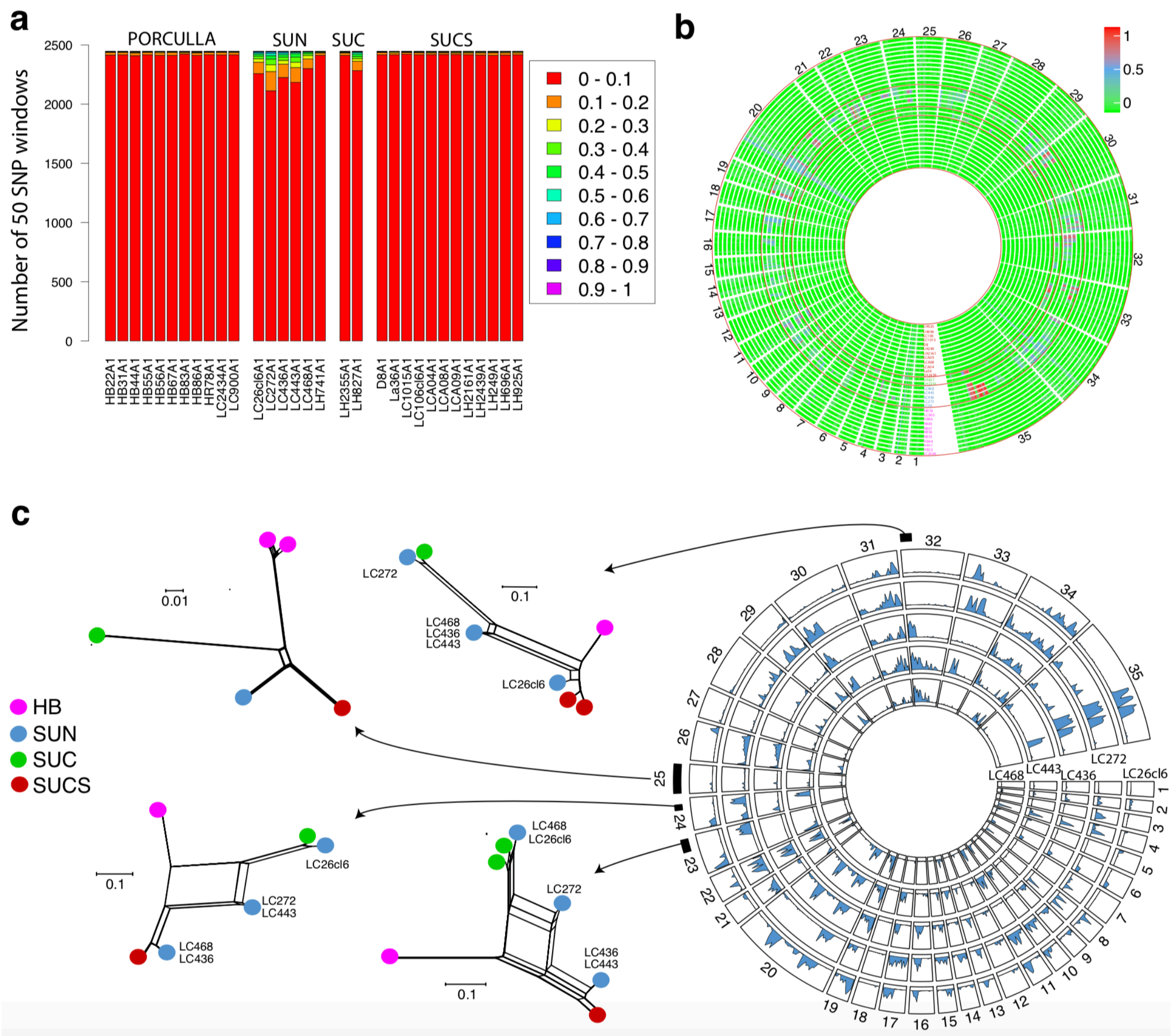

**Supplementary Figure 9**. **(a-b)** Proportion of heterozygous sites in 50 SNP windows for each *L. peruviana* genome revealed that all SUN isolates and SUC isolate LH827 contained genomic windows with elevated proportions of heterozygous sites compared to the other *L. peruviana* isolates. **(c)** Phylogenetic networks of SNPs in several such heterozygous regions revealed that these isolates were of mixed ancestry. For instance, isolate LC272 contained elevated proportions of heterozygous sites in chromosome 23, and this isolate clustered between the SUC and the SUCS groups. In the same genomic region, SUN isolates LC436 and LC443 were largely homozygous and clustered entirely with the SUCS population, suggesting introgressive hybridization between SUCS and SUN. Similar patterns are shown for chromosome 24 and 32. For chromosome 25 that was largely homozygous in all isolates, network analyses showed a similar population structure as observed based on genome-wide SNPs (Supp. Fig. 6, 7b).

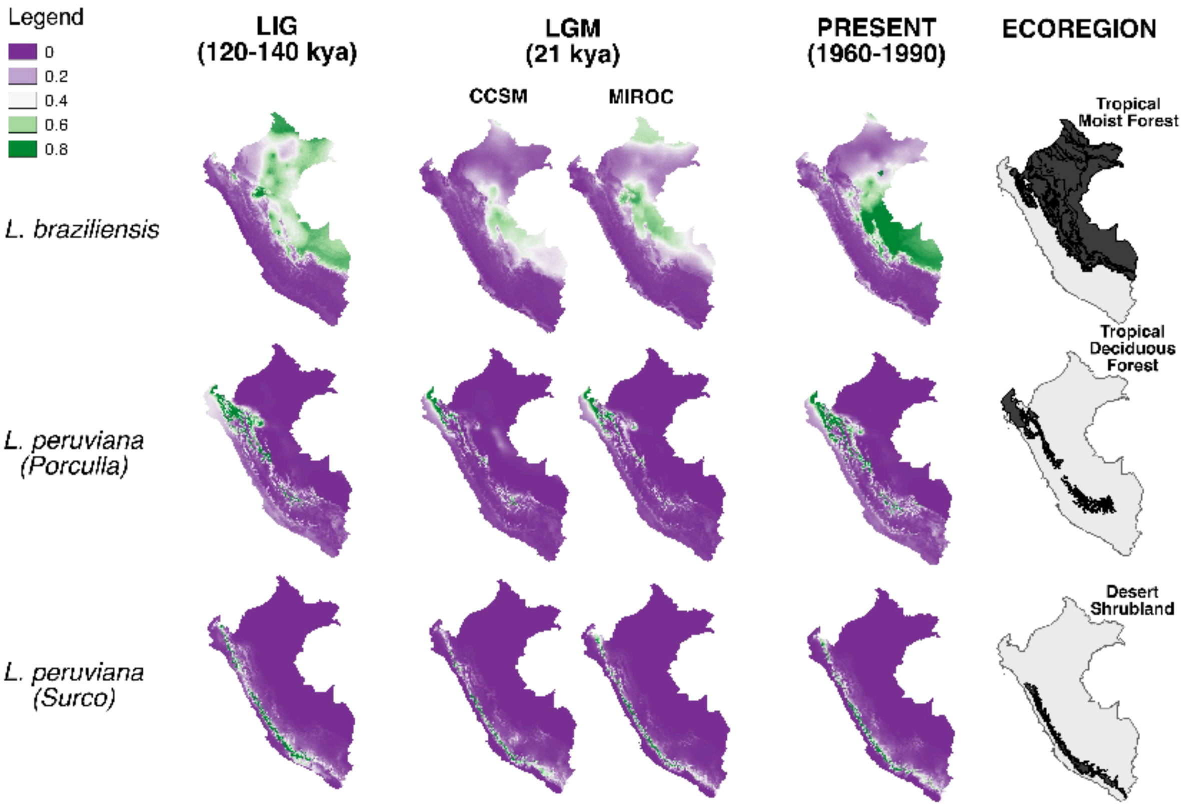

**Supplementary Figure 10**. Tests for range expansion, contraction or shift using ecological niche modeling. Geographic maps of Peru depicting the distribution probability for each of the three major Leishmania groups for the present, the last interglacial (LIG) and the last glacial maximum (LGM) under different climate models: the Community Climate System Model25 (CCSM) and the Model for Interdisciplinary Research on Climate26 (MIROC).

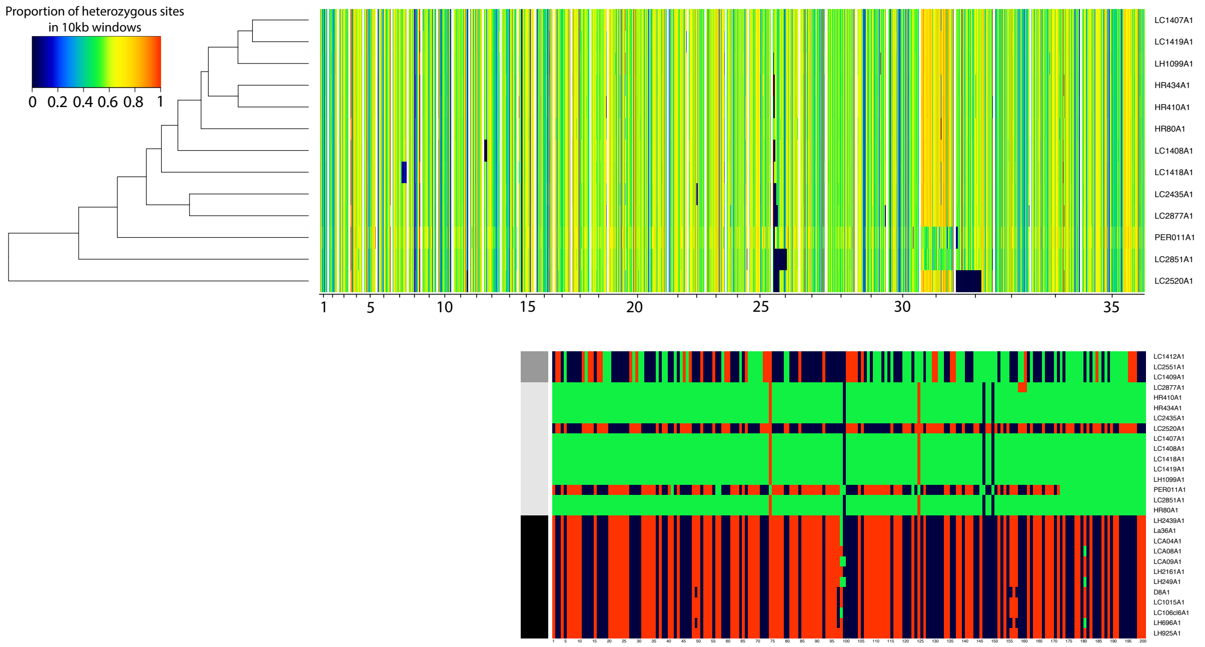

**Supplementary Figure 11.** Heatmap on top shows the proportion of heterozygous sites in 10kb windows across the 35 major chromosomes (x-axis) for the 13 hybrid *L. peruviana* x *L. braziliensis* isolates. Heatmap below shows the allelic profiles of the first 86.5kb of chromosome 32 in the 13 hybrid isolates (light grey boxes), their putative *L. peruviana* parent group (black boxes) and their putative *L. braziliensis* parent group (dark grey boxes). For each SNP identified within this stretch, alleles were coloured dark blue if homozygous for the reference allele, dark red if homozygous for the alternate allele and green if heterozygous. Notice the two homozygous stretches in isolate LC2520 (homozygous for the *L. braziliensis* alternate alleles) and PER011(homozygous *L. peruviana* alternate alleles).

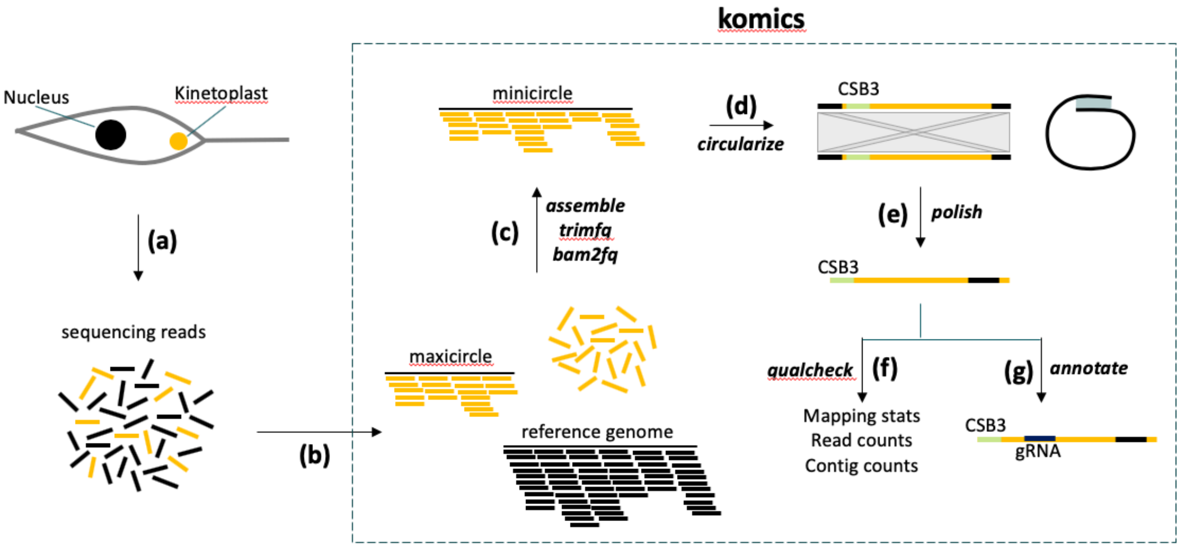

**Supplementary Figure 12.** The KOMICS pipeline as implemented in a python package, showing the different modules at each step (see text for details). **(a)** Whole cellular DNA extractions are used for paired-end whole-genome sequencing with Illumina. **(b)** The reads are aligned to the reference genome containing the 35 major chromosomes and a complete maxicircle sequence. **(c)** Unaligned reads are extracted from the BAM files, quality trimmed and used for *de novo* assembly. **(d)** Contigs containing the CSB3 reads are extracted and circularized using a BLAST approach. **(e)** Circularized minicircle sequences were further polished by putting the CSB3-mer at the start of the minicircle sequence. **(f)** Unaligned reads from step **(b)** are mapped to the final sets of minicircles to calculate mapping statistics. **(g)** Guide RNA genes are predicted by aligning minicircles to edited maxicircle sequences (this step is not implemented in KOMICS).

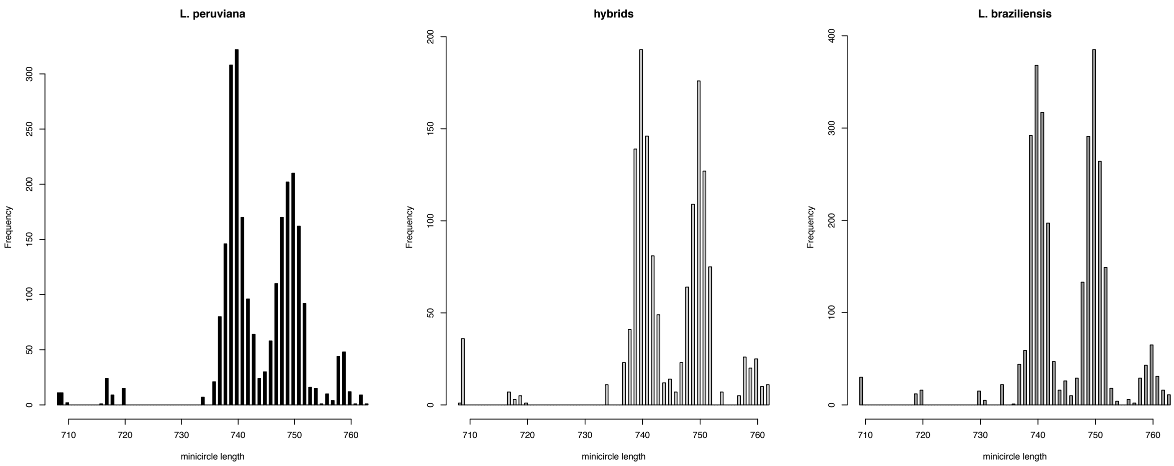

**Supplementary Figure 13.** Minicircle length distribution for each species and their hybrids.

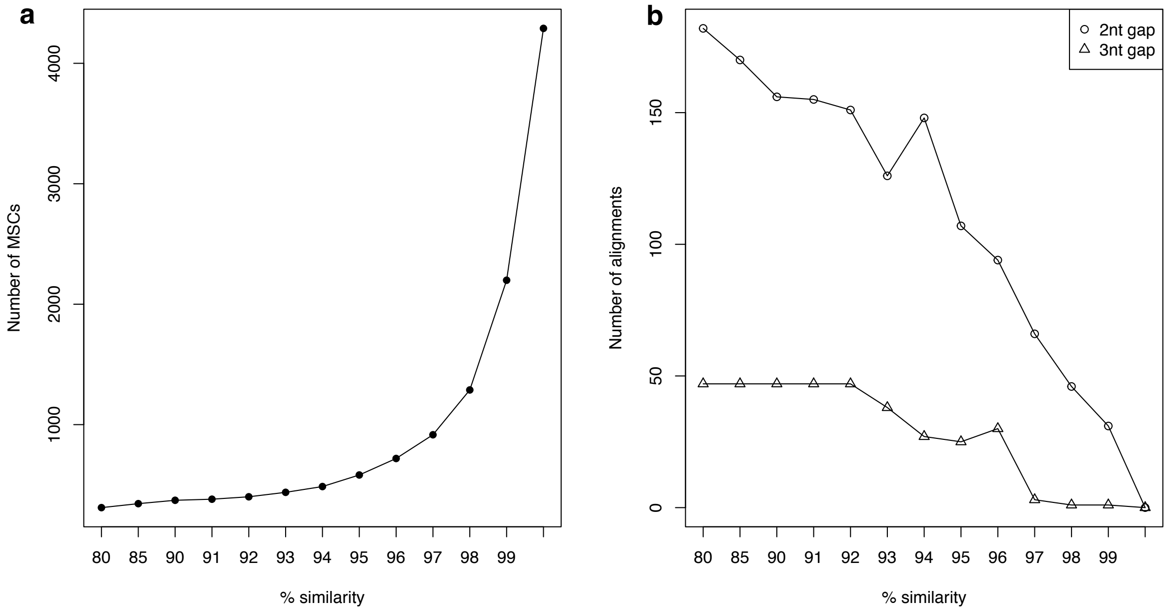

**Supplementary Figure 14. (a)** Results of clustering analyses whereby minicircle sequences were grouped into minicircle sequence classes (MSC) (y-axis) based on percent similarity (x-axis). **(b)** Number of alignments with a 2nt and 3nt gap during the clustering analyses.

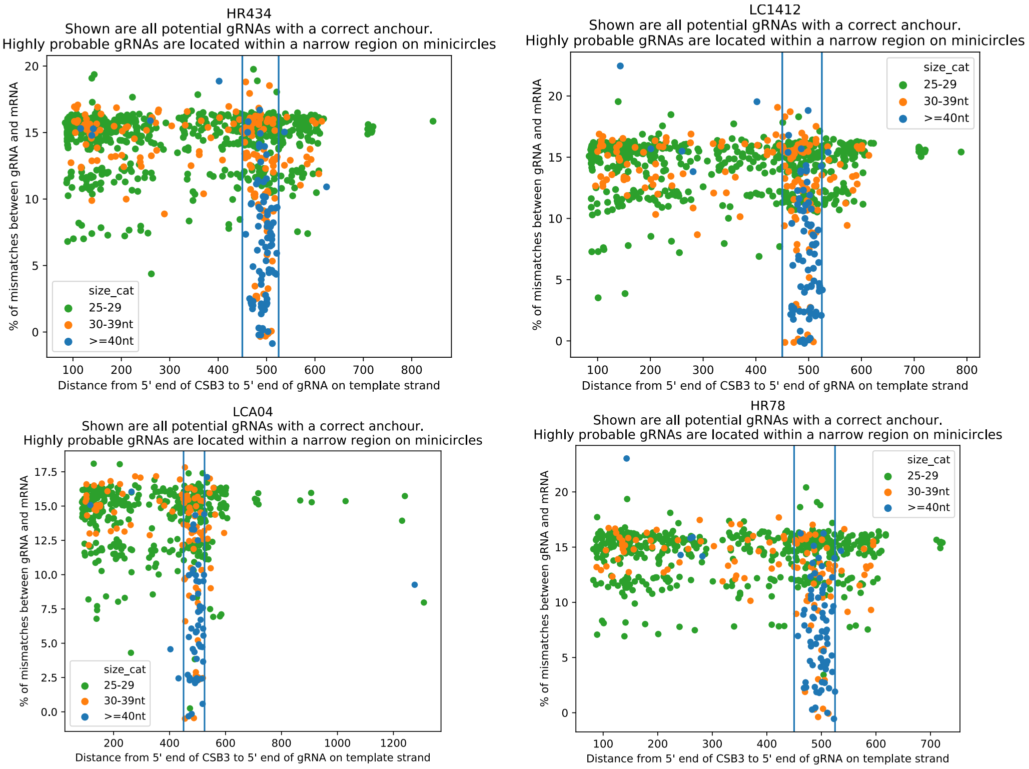

**Supplementary Figure 15.** Positions of predicted gRNA's relative to the CSB3 fragment on each minicircle. Only gRNA's within 500bp from the CSB3 are kept for further analyses.

### SUPPLEMENTARY TABLES

**Table S1. List of 68 isolates used for whole-genome sequencing.**
Note that - for the purpuses of minicircle sequence analyses - four isolates were re-sequenced after a phenol/chloroform DNA extraction, resulting in a total of 72 whole genomes. For anonymity reasons, details on sampling locations were omitted from the table, but are availabe upon request.
The World Health Organization (WHO) code is as follows: host (M for Mammalia, HOM = Homo sapiens & CAN = Canis familiaris; I for Insecta; AYA = Lutzomyia ayacuchensis)/country(BO = Bolivia; PE = Peru)/year of isolation/name of strain.

| WHO code | Sample Code | Species | Year of isolation | Country | Region | DNA extraction |
|---|---|---|---|---|---|---|
| **MHOM/PE/02/LH2210(PER010)** | PER010A1 | *L. braziliensis* | 2002 | Peru | Cajamarca | commercial kit |
| **MHOM/PE/91/LC1565** | LC1565A1 | *L. braziliensis* | 1991 | Peru | Cusco | commercial kit |
| **MHOM/PE/01/LH2217(PER012)** | PER012A1 | *L. braziliensis* | 2001 | Peru | Cusco | commercial kit |
| **MHOM/PE/02/LH2287(PER065)** | PER065A1 | *L. braziliensis* | 2002 | Peru | Cusco | commercial kit |
| **MHOM/PE/91/LC1409** | LC1409A1 | *L. braziliensis* | 1991 | Peru | Huánuco | commercial kit |
| **MHOM/PE/91/LC1412** | LC1412A1 | *L. braziliensis* | 1991 | Peru | Huánuco | commercial kit |
| **MCAN/PE/94/LC2412** | LC2421A1 | *L. braziliensis* | 1994 | Peru | Huánuco | commercial kit |
| **MHOM/PE/94/LC2452** | LC2452A1 | *L. braziliensis* | 1994 | Peru | Huánuco | commercial kit |
| **MHOM/PE/94/LC2551** | LC2551A1 | *L. braziliensis* | 1994 | Peru | Huánuco | commercial kit |
| **MHOM/PE/95/LC2873** | LC2873A1 | *L. braziliensis* | 1995 | Peru | Huánuco | commercial kit |
| **MHOM/PE/02/LH2224(PER016)** | PER016A1 | *L. braziliensis* | 2002 | Peru | Huánuco | commercial kit |
| **MHOM/PE/02/LH2356(PER094)** | PER094A1 | *L. braziliensis* | 2002 | Peru | Huánuco | commercial kit |
| **MHOM/PE/01/LH2033(PER014)** | PER014A1 | *L. braziliensis* | 2001 | Peru | Junín | commercial kit |
| **MHOM/PE/01/LH2182(PER005)** | PER005A1 | *L. braziliensis* | 2001 | Peru | Loreto | commercial kit |
| **MHOM/PE/03/PER201** | PER201A1 | *L. braziliensis* | 2003 | Peru | Loreto | commercial kit |
| **MHOM/PE/01/LH2162(PER002)** | PER002A1 | *L. braziliensis* | 2001 | Peru | Madre de Dios | commercial kit |
| **MHOM/PE/02/LH2330(PER069)** | PER069A1 | *L. braziliensis* | 2002 | Peru | Madre de Dios | commercial kit |
| **MHOM/PE/91/LC1412** | LC1412C3 | *L. braziliensis* | 1991 | Peru | Huánuco | phenol/chloroform |
| **MHOM/PE/03/PER260** | PER260A1 | *L. braziliensis* | 2003 | Peru | Madre de Dios | commercial kit |
| **MHOM/BO/94/CUM29** | CUM29A1 | *L. braziliensis* | 1994 | Bolivia | Parque Isiboro | commercial kit |
| **MHOM/PE/02/LH2332(PER086)** | PER086A1 | *L. braziliensis* | 2002 | Peru | Pasco | commercial kit |
| **MHOM/BR/2015/RO393** | RO393A1 | *L. braziliensis* | 2015 | Brazil | Rondônia | commercial kit |
| **MHOM/PE/90/LH825** | LH825B2 | *L. braziliensis* | 1990 | Peru | Ucayali | commercial kit |
| **MHOM/PE/03/PER215** | PER215A1 | *L. braziliensis* | 2003 | Peru | Ucayali | commercial kit |
| **MHOM/CO/83/REST417** | REST417A1 | *L. panamensis* | 1983 | Colombia | - | phenol/chloroform |
| **MCAN/PE/95/HR434** | HR434C3 | hybrid | 1995 | Peru | Huánuco | phenol/chloroform |
| **MCAN/PE/95/HR410** | HR410A1 | hybrid | 1995 | Peru | Huánuco | commercial kit |
| **MCAN/PE/95/HR434** | HR434A1 | hybrid | 1995 | Peru | Huánuco | commercial kit |
| **MCAN/PE/95/HR80** | HR80A1 | hybrid | 1995 | Peru | Huánuco | commercial kit |
| **MHOM/PE/91/LC1407** | LC1407A1 | hybrid | 1991 | Peru | Huánuco | commercial kit |
| **MHOM/PE/91/LC1408** | LC1408A1 | hybrid | 1991 | Peru | Huánuco | commercial kit |
| **MHOM/PE/91/LC1418** | LC1418A1 | hybrid | 1991 | Peru | Huánuco | commercial kit |
| **MHOM/PE/91/LC1419** | LC1419A1 | hybrid | 1991 | Peru | Huánuco | commercial kit |
| **MHOM/PE/94/LC2435** | LC2435A1 | hybrid | 1994 | Peru | Huánuco | commercial kit |
| **MHOM/PE/94/LC2520** | LC2520A1 | hybrid | 1994 | Peru | Huánuco | commercial kit |

| | | | | | | |
|---|---|---|---|---|---|---|
| **MHOM/PE/95/LC2851** | LC2851A1 | hybrid | 1995 | Peru | Huánuco | commercial kit |
| **MHOM/PE/95/LC2877** | LC2877A1 | hybrid | 1995 | Peru | Huánuco | commercial kit |
| **MHOM/PE/91/LH1099** | LH1099A1 | hybrid | 1991 | Peru | Huánuco | commercial kit |
| **MHOM/PE/02/LH2215(PER011)** | PER011A1 | hybrid | 2002 | Peru | Huánuco | commercial kit |
| **MHOM/PE/03/LH2439(PER126)** | LH2439A1 | *L. peruviana* | 2003 | Peru | - | commercial kit |
| **MHOM/PE/84/LC26cl6** | LC26cl6A1 | *L. peruviana* | 1984 | Peru | Ancash | commercial kit |
| **MHOM/PE/88/LC272** | LC272A1 | *L. peruviana* | 1988 | Peru | Ancash | commercial kit |
| **MHOM/PE/90/LC436** | LC436A1 | *L. peruviana* | 1990 | Peru | Ancash | commercial kit |
| **MHOM/PE/90/LC443** | LC443A1 | *L. peruviana* | 1990 | Peru | Ancash | commercial kit |
| **MHOM/PE/90/LC468** | LC468A1 | *L. peruviana* | 1990 | Peru | Ancash | commercial kit |
| **MHOM/PE/89/LH741** | LH741A1 | *L. peruviana* | 1989 | Peru | Ancash | commercial kit |
| **IAYA/PE/90/La36** | La36A1 | *L. peruviana* | 1990 | Peru | Ayacucho | commercial kit |
| **MHOM/PE/90/LCA04** | LCA04A1 | *L. peruviana* | 1990 | Peru | Ayacucho | commercial kit |
| **MHOM/PE/90/LCA08** | LCA08A1 | *L. peruviana* | 1990 | Peru | Ayacucho | commercial kit |
| **MHOM/PE/90/LCA09** | LCA09A1 | *L. peruviana* | 1990 | Peru | Ayacucho | commercial kit |
| **MHOM/PE/01/LH2161(PER001)** | LH2161A1 | *L. peruviana* | 2001 | Peru | Ayacucho | commercial kit |
| **MHOM/PE/90/LH249** | LH249A1 | *L. peruviana* | 1990 | Peru | Ayacucho | commercial kit |
| **MCAN/PE/95/HR78** | HR78C3 | *L. peruviana* | 1995 | Peru | Quera | phenol/chloroform |
| **MCAN/PE/95/HR78** | HR78A1 | *L. peruviana* | 1995 | Peru | Huánuco | commercial kit |
| **MHOM/PE/94/LC2434** | LC2434A1 | *L. peruviana* | 1994 | Peru | Huánuco | commercial kit |
| **MCAN/PE/76/D8(LV608)** | D8A1 | *L. peruviana* | 1976 | Peru | Lima | commercial kit |
| **MHOM/PE/91/LC1015** | LC1015A1 | *L. peruviana* | 1991 | Peru | Lima | commercial kit |
| **MHOM/PE/90/LCA04** | LCA04B2 | *L. peruviana* | 1990 | Peru | Ayacucho | phenol/chloroform |
| **MHOM/PE/85/LC106cl6** | LC106cl6A1 | *L. peruviana* | 1985 | Peru | Lima | commercial kit |
| **MHOM/PE/02/LH2355(PER087)** | LH2355A1 | *L. peruviana* | 2002 | Peru | Lima | commercial kit |
| **MHOM/PE/89/LH696** | LH696A1 | *L. peruviana* | 1989 | Peru | Lima | commercial kit |
| **MHOM/PE/90/LH827** | LH827A1 | *L. peruviana* | 1990 | Peru | Lima | commercial kit |
| **MHOM/PE/90/LH925** | LH925A1 | *L. peruviana* | 1990 | Peru | Lima | commercial kit |
| **MHOM/PE/90/HB22** | HB22A1 | *L. peruviana* | 1990 | Peru | Piura | commercial kit |
| **MHOM/PE/90/HB31** | HB31A1 | *L. peruviana* | 1990 | Peru | Piura | commercial kit |
| **MHOM/PE/90/HB44** | HB44A1 | *L. peruviana* | 1990 | Peru | Piura | commercial kit |
| **MHOM/PE/90/HB55** | HB55A1 | *L. peruviana* | 1990 | Peru | Piura | commercial kit |
| **MHOM/PE/90/HB56** | HB56A1 | *L. peruviana* | 1990 | Peru | Piura | commercial kit |
| **MHOM/PE/90/HB67** | HB67A1 | *L. peruviana* | 1990 | Peru | Piura | commercial kit |
| **MHOM/PE/90/HB83** | HB83A1 | *L. peruviana* | 1990 | Peru | Piura | commercial kit |
| **MHOM/PE/90/HB86** | HB86A1 | *L. peruviana* | 1990 | Peru | Piura | commercial kit |
| **MHOM/PE/89/LC900** | LC900A1 | *L. peruviana* | 1989 | Peru | Piura | commercial kit |

**Supplementary Table 2:** Deleterious mutations (SNPs and INDELs) that were fixed in *L. peruviana* and absent in *L. braziliensis.* All INDELs occured in hypothetical proteins.

| TYPE OF MUTATION | CHROMOSOME | POSITION | REFERENCE ALLELE | ALTERNATE ALLEL | EFFECT | PRODUCT | ORTHOLOGOUS GROUP | GENE ID | NEW GENE ID |
|---|---|---|---|---|---|---|---|---|---|
| SNP | LbrM_17_v4_49 | 664535 | C | T | stop gained | ion transport protein, putative | ORTHOMCL5645 | LbrM.17.1590 | LbrM2904_000278800.1 |
| SNP | LbrM_20_v4_69 | 395387 | T | C | start lost | translation elongation factor 1-beta (eEF1B beta 2) | NA | NA | LbrM2904_000325200.1 |
| DELETION | LbrM_23_v4_56 | 278811 | TG | T | frameshift | hypothetical | ORTHOMCL6525 | LbrM.11.0220 | LbrM2904_000435600.1 |
| SNP | LbrM_23_v4_56 | 257291 | A | G | start lost | kinesin-C | ORTHOMCL4417 | LbrM.23.0710 | LbrM2904_000435000.1 |
| INSERTION | LbrM_25_v4_58 | 515739 | T | TCA | frameshift | hypothetical | ORTHOMCL3965 | LbrM.25.1270 | LbrM2904_000487800.1 |
| INSERTION | LbrM_28_v4_62 | 1109098 | G | GC | frameshift | hypothetical | ORTHOMCL3110 | LbrM.28.2900 | LbrM2904_000586800.1 |
| SNP | LbrM_34_v4_67 | 1231005 | A | G | start lost | class I transcription factor A, subunit 2, putative | ORTHOMCL1229 | LbrM.34.3060 | LbrM2904_000809700.1 |
| DELETION | LbrM_35_v4_68 | 842557 | GTC | G | frameshift | hypothetical | NA | NA | LbrM2904_000857400.1 |
| SNP | LbrM_35_v4_68 | 915726 | C | T | stop gained | hypothetical | ORTHOMCL817 | LbrM.35.2530 | LbrM2904_000859300.1 |

**Supplementary Table 3:** *F* IS as estimated per polymorphic SNP site in three groups of parasites representing the three major *Leishmania populations* in Peru.

| Population | *L. braziliensis* | *L. peruviana* Porculla | *L. peruviana* Surco |
|---|---|---|---|
| Country | Peru | Peru | Peru |
| Region | Huánuco | Piura | Ayacucho |
| Year | 1991-1995 | 1989-1990 | 1990 |
| N isolates | 6 | 4 | 5 |
| N polymorphic sites | 162,204 | 1,877 | 1,426 |
| Proportion of loci with -0.1 > FIS < 0.1 | 40% | 3,56% | 0% |
| Proportion of loci with FIS < -0.1 | 44% | 90% | 93% |
| Proportion of loci with FIS = -1 | 2,38% | 42% | 48% |
| Fis median | -0,09 | -0,6 | -0,67 |
| Fis mean | -0,11 | -0,54 | -0,54 |

**Supplementary Table 4:** Number of SNPs differing at one (below diagonal) or both (above diagonal) alleles. Hybrids from the Nolder et al. (2009) are indicated by their respective zymodeme between brackets (LON2018-221)

| Column1 | LC2877A1 (LON221) | HR410A1 (LON219) | HR434A1 (LON218) | LC2435A1 (LON220) | HR80A1 (LON218) | LC2520A1 | LC1407A1 | LC1408A1 | LC1418A1 | LC1419A1 | LH1099A1 | LC2851A1 | PER011A1 |
|---|---|---|---|---|---|---|---|---|---|---|---|---|---|
| **LC2877A1 (LON221)** | | 0 | 0 | 0 | 1 | 4 | 1 | 1 | 2 | 0 | 0 | 2 | 3 |
| **HR410A1 (LON219)** | 1036 | | 0 | 1 | 0 | 12 | 1 | 0 | 1 | 0 | 0 | 0 | 0 |
| **HR434A1 (LON218)** | 1042 | 322 | | 1 | 0 | 12 | 1 | 0 | 1 | 0 | 0 | 2 | 0 |
| **LC2435A1 (LON220)** | 884 | 662 | 668 | | 0 | 3 | 1 | 0 | 1 | 0 | 0 | 1 | 0 |
| **HR80A1 (LON218)** | 1255 | 825 | 841 | 1053 | | 3 | 1 | 0 | 2 | 0 | 0 | 4 | 0 |
| **LC2520A1** | 3928 | 4088 | 4106 | 3962 | 4391 | | 5 | 4 | 6 | 3 | 3 | 4 | 167 |
| **LC1407A1** | 1316 | 888 | 906 | 1144 | 569 | 4462 | | 1 | 2 | 1 | 1 | 1 | 1 |
| **LC1408A1** | 1235 | 875 | 923 | 1061 | 1036 | 4381 | 869 | | 2 | 0 | 0 | 1 | 2 |
| **LC1418A1** | 1598 | 1110 | 1144 | 1356 | 1103 | 4698 | 946 | 1217 | | 1 | 1 | 2 | 1 |
| **LC1419A1** | 1264 | 834 | 874 | 1084 | 527 | 4422 | 326 | 835 | 916 | | 0 | 0 | 0 |
| **LH1099A1** | 1255 | 767 | 803 | 1029 | 790 | 4359 | 615 | 884 | 825 | 557 | | 0 | 0 |
| **LC2851A1** | 3578 | 3792 | 3784 | 3656 | 4007 | 6426 | 4110 | 4025 | 4368 | 4070 | 4051 | | 0 |
| **PER011A1** | 6293 | 5851 | 5857 | 6111 | 6042 | 9103 | 5919 | 5976 | 6125 | 5867 | 5792 | 9111 | |

**Supplementary Table 5:** Mapping statistics as outputted by the KOMICS pipeline following assembly and circularization of minicircle sequence classes per isolate.

| Isolate | Median genome-wide read depth | Number of reads | Number of mapped reads | Proportion of mapped reads | Number of reads w/ MQ>20 | Proportion of mapped reads w/ MQ>20 | Number of proper pairs | Proportion of mapped proper pairs | Number of CSB3 reads | Number of mapped CSB3 reads | Proportion of mapped CSB3 reads | Number of perfectly matched CSB3 reads | Proportion of perfectly matched CSB3 read | Number of proper paired CSB3 reads | Proportion of properly paried CSB3 reads | Number of minicircle sequence classes |
|---|---|---|---|---|---|---|---|---|---|---|---|---|---|---|---|---|
| **MEAN** | | **1938903** | - | **80%** | - | **93%** | - | **95%** | - | - | **95%** | - | **89%** | - | **90%** | - |
| **MEDIAN** | | **1704236** | - | **81%** | - | **93%** | - | **95%** | - | - | **95%** | - | **91%** | - | **91%** | - |
| **MIN** | | **437178** | - | **36%** | - | **89%** | - | **91%** | - | - | **91%** | - | **81%** | - | **83%** | - |
| **MAX** | | **6022478** | - | **94%** | - | **95%** | - | **97%** | - | - | **98%** | - | **96%** | - | **95%** | - |
| CUM29A1 | 63 | 2563732 | 2249534 | 88% | 2075048 | 92% | 2151222 | 96% | 236609 | 222890 | 94% | 208320 | 88% | 213562 | 90% | 274 |
| D8A1 | 64 | 856842 | 703178 | 82% | 647624 | 92% | 662316 | 94% | 80883 | 77885 | 96% | 73829 | 91% | 74201 | 92% | 87 |
| HB22A1 | 87 | 3821476 | 3369185 | 88% | 3133343 | 93% | 3229178 | 96% | 427851 | 410095 | 96% | 393514 | 92% | 395799 | 93% | 109 |
| HB31A1 | 69 | 4160628 | 3632897 | 87% | 3323128 | 91% | 3442252 | 95% | 476648 | 448164 | 94% | 422812 | 89% | 428076 | 90% | 111 |
| HB44A1 | 99 | 4210622 | 3602197 | 86% | 3330085 | 92% | 3425858 | 95% | 465971 | 442432 | 95% | 423098 | 91% | 417804 | 90% | 111 |
| HB55A1 | 89 | 1486450 | 1176572 | 79% | 1083480 | 92% | 1117762 | 95% | 152011 | 145561 | 96% | 140548 | 92% | 139356 | 92% | 107 |
| HB56A1 | 81 | 3110238 | 2752880 | 89% | 2554937 | 93% | 2641650 | 96% | 339739 | 327589 | 96% | 312080 | 92% | 315044 | 93% | 110 |
| HB67A1 | 61 | 4427034 | 1587847 | 36% | 1464123 | 92% | 1523672 | 96% | 188816 | 181984 | 96% | 173117 | 92% | 174866 | 93% | 111 |
| HB83A1 | 77 | 3415300 | 3017341 | 88% | 2805209 | 93% | 2870352 | 95% | 364649 | 345222 | 95% | 329588 | 90% | 332356 | 91% | 109 |
| HB86A1 | 80 | 1877216 | 1562713 | 83% | 1439785 | 92% | 1487988 | 95% | 194191 | 181877 | 94% | 172605 | 89% | 173945 | 90% | 101 |
| HR410A1 | 55 | 505250 | 343951 | 68% | 321899 | 94% | 315576 | 92% | 68370 | 62068 | 91% | 56256 | 82% | 57726 | 84% | 123 |
| HR434A1 | 60 | 609148 | 458471 | 75% | 437281 | 95% | 435640 | 95% | 83980 | 78668 | 94% | 72129 | 86% | 75318 | 90% | 112 |
| HR78A1 | 59 | 437178 | 275903 | 63% | 262137 | 95% | 257710 | 93% | 48974 | 45396 | 93% | 41579 | 85% | 42662 | 87% | 104 |
| HR80A1 | 53 | 765524 | 604022 | 79% | 566868 | 94% | 560856 | 93% | 122025 | 113173 | 93% | 103156 | 85% | 106493 | 87% | 133 |
| La36A1 | 87 | 972144 | 784008 | 81% | 723533 | 92% | 741524 | 95% | 97859 | 93823 | 96% | 89897 | 92% | 90082 | 92% | 85 |
| LC1015A1 | 62 | 784688 | 634434 | 81% | 592461 | 93% | 607436 | 96% | 79594 | 76060 | 96% | 72634 | 91% | 73000 | 92% | 78 |
| LC106cl6A1 | 88 | 2652184 | 2384321 | 90% | 2209944 | 93% | 2255728 | 95% | 307178 | 293576 | 96% | 281537 | 92% | 283220 | 92% | 88 |
| LC1407A1 | 76 | 637664 | 430846 | 68% | 389924 | 91% | 399328 | 93% | 59160 | 54446 | 92% | 51323 | 87% | 49257 | 83% | 132 |
| LC1408A1 | 87 | 1389504 | 1133765 | 82% | 1037637 | 92% | 1080672 | 95% | 145306 | 138621 | 95% | 131482 | 90% | 132467 | 91% | 150 |
| LC1418A1 | 87 | 1042116 | 818559 | 79% | 748063 | 91% | 771514 | 94% | 107806 | 103301 | 96% | 98702 | 92% | 97749 | 91% | 148 |
| LC1419A1 | 105 | 1390264 | 1116626 | 80% | 1021255 | 91% | 1057354 | 95% | 147463 | 141159 | 96% | 134893 | 91% | 134836 | 91% | 138 |
| LC2421A1 | 62 | 817308 | 621914 | 76% | 579007 | 93% | 572132 | 92% | 119664 | 111040 | 93% | 100744 | 84% | 102459 | 86% | 176 |
| LC2434A1 | 50 | 459972 | 322401 | 70% | 304542 | 94% | 300432 | 93% | 63615 | 59030 | 93% | 53576 | 84% | 55037 | 87% | 90 |
| LC2435A1 | 72 | 489270 | 264253 | 54% | 249994 | 95% | 245250 | 93% | 50960 | 46813 | 92% | 42845 | 84% | 43978 | 86% | 123 |
| LC2452A1 | 58 | 670500 | 405465 | 60% | 380528 | 94% | 373244 | 92% | 81474 | 74038 | 91% | 66332 | 81% | 69264 | 85% | 174 |
| LC2520A1 | 65 | 527630 | 335432 | 64% | 317116 | 95% | 315510 | 94% | 61813 | 57405 | 93% | 52539 | 85% | 54344 | 88% | 114 |
| LC2551A1 | 59 | 870492 | 648749 | 75% | 612130 | 94% | 600398 | 93% | 130842 | 120029 | 92% | 107144 | 82% | 112544 | 86% | 108 |
| LC26cl6A1 | 82 | 835740 | 698582 | 84% | 652667 | 93% | 676538 | 97% | 87046 | 85314 | 98% | 82106 | 94% | 82846 | 95% | 99 |
| LC272A1 | 94 | 2547878 | 2266474 | 89% | 2117227 | 93% | 2188840 | 97% | 287367 | 276602 | 96% | 263646 | 92% | 268773 | 94% | 100 |
| LC2851A1 | 53 | 479088 | 333703 | 70% | 316633 | 95% | 307554 | 92% | 64979 | 60087 | 92% | 55083 | 85% | 55763 | 86% | 117 |
| LC2873A1 | 63 | 742998 | 544601 | 73% | 506953 | 93% | 497002 | 91% | 110518 | 100649 | 91% | 89391 | 81% | 93009 | 84% | 251 |
| LC2877A1 | 50 | 703758 | 547038 | 78% | 513484 | 94% | 508914 | 93% | 111153 | 102061 | 92% | 92313 | 83% | 96321 | 87% | 106 |
| LC436A1 | 123 | 2027388 | 1632141 | 81% | 1508194 | 92% | 1542344 | 94% | 203986 | 198548 | 97% | 188811 | 93% | 185061 | 91% | 103 |
| LC443A1 | 80 | 2020436 | 1701260 | 84% | 1568366 | 92% | 1605280 | 94% | 230426 | 218803 | 95% | 209185 | 91% | 203902 | 88% | 99 |
| LC468A1 | 93 | 1835794 | 1624660 | 88% | 1505595 | 93% | 1562226 | 96% | 207732 | 200047 | 96% | 190272 | 92% | 193685 | 93% | 106 |
| LC900A1 | 89 | 3353872 | 3022193 | 90% | 2786692 | 92% | 2874660 | 95% | 358752 | 343569 | 96% | 329472 | 92% | 331039 | 92% | 110 |
| LCA04A1 | 91 | 2211264 | 1975696 | 89% | 1839578 | 93% | 1883632 | 95% | 241437 | 233534 | 97% | 221397 | 92% | 225952 | 94% | 88 |
| LCA08A1 | 94 | 531410 | 399713 | 75% | 372246 | 93% | 383864 | 96% | 50276 | 48816 | 97% | 46497 | 92% | 47252 | 94% | 85 |
| LCA09A1 | 100 | 750612 | 592146 | 79% | 555424 | 94% | 570646 | 96% | 72186 | 70241 | 97% | 67437 | 93% | 67900 | 94% | 84 |
| LH1099A1 | 89 | 774338 | 557781 | 72% | 505573 | 91% | 523360 | 94% | 77308 | 73924 | 96% | 70982 | 92% | 69621 | 90% | 130 |
| LH2161A1 | 94 | 2060770 | 1631760 | 79% | 1512947 | 93% | 1548654 | 95% | 207682 | 195460 | 94% | 182858 | 88% | 185322 | 89% | 87 |
| LH2355A1 | 110 | 1565798 | 1215102 | 78% | 1135240 | 93% | 1168720 | 96% | 151139 | 147228 | 97% | 142834 | 95% | 141614 | 94% | 92 |
| LH2439A1 | 87 | 961024 | 809411 | 84% | 755121 | 93% | 771252 | 95% | 105302 | 99516 | 95% | 92737 | 88% | 96146 | 91% | 85 |
| LH249A1 | 79 | 1235982 | 1037959 | 84% | 971023 | 94% | 994734 | 96% | 125723 | 122267 | 97% | 116304 | 93% | 118051 | 94% | 80 |
| LH696A1 | 85 | 1704236 | 1502320 | 88% | 1400521 | 93% | 1424218 | 95% | 174990 | 169270 | 97% | 162718 | 93% | 162827 | 93% | 89 |
| LH741A1 | 73 | 2062296 | 1843172 | 89% | 1717217 | 93% | 1781062 | 97% | 223733 | 214198 | 96% | 203306 | 91% | 206322 | 92% | 81 |
| LH825B2 | 56 | 480114 | 324564 | 68% | 302314 | 93% | 300340 | 93% | 64473 | 59097 | 92% | 52734 | 82% | 55312 | 86% | 293 |
| LH827A1 | 82 | 2085308 | 1874883 | 90% | 1753588 | 94% | 1809228 | 96% | 237857 | 231093 | 97% | 219899 | 92% | 224262 | 94% | 103 |
| LH925A1 | 73 | 1773630 | 1524147 | 86% | 1415896 | 93% | 1419932 | 93% | 179503 | 169877 | 95% | 163464 | 91% | 159192 | 89% | 81 |
| PER002A1 | 59 | 2794564 | 2068061 | 74% | 1838627 | 89% | 1937296 | 94% | 293638 | 271941 | 93% | 243261 | 83% | 246800 | 84% | 318 |
| PER005A1 | 50 | 1857180 | 1486084 | 80% | 1376300 | 93% | 1427564 | 96% | 176911 | 170968 | 97% | 160581 | 91% | 163085 | 92% | 213 |
| PER010A1 | 70 | 1203276 | 969626 | 81% | 878669 | 91% | 916082 | 94% | 120775 | 113962 | 94% | 104299 | 86% | 106106 | 88% | 333 |
| PER011A1 | 53 | 4611804 | 4260033 | 92% | 3982221 | 93% | 4104410 | 96% | 473537 | 463141 | 98% | 442094 | 93% | 445026 | 94% | 233 |
| PER012A1 | 57 | 4015010 | 3567576 | 89% | 3251468 | 91% | 3400194 | 95% | 424394 | 400597 | 94% | 369337 | 87% | 376795 | 89% | 303 |
| PER014A1 | 68 | 2791912 | 2553819 | 91% | 2374264 | 93% | 2448794 | 96% | 282118 | 267541 | 95% | 253824 | 90% | 257068 | 91% | 155 |
| PER065A1 | 62 | 4872468 | 4234412 | 87% | 3897910 | 92% | 4004884 | 95% | 519873 | 487354 | 94% | 451079 | 87% | 443694 | 85% | 188 |
| PER069A1 | 62 | 2470734 | 2228883 | 90% | 2048915 | 92% | 2169834 | 97% | 245228 | 236180 | 96% | 223440 | 91% | 231091 | 94% | 123 |
| PER086A1 | 62 | 4431772 | 2625497 | 59% | 2417373 | 92% | 2504756 | 95% | 312722 | 298344 | 95% | 280767 | 90% | 283858 | 91% | 208 |
| PER094A1 | 86 | 6022478 | 4793749 | 80% | 4394258 | 92% | 4621366 | 96% | 567178 | 538391 | 95% | 495673 | 87% | 517226 | 91% | 246 |
| PER201A1 | 58 | 2146594 | 1986552 | 93% | 1846268 | 93% | 1927370 | 97% | 228326 | 220160 | 96% | 208002 | 91% | 210853 | 92% | 184 |
| PER215A1 | 60 | 2744252 | 2573124 | 94% | 2423950 | 94% | 2481362 | 96% | 292068 | 287177 | 98% | 280185 | 96% | 274126 | 94% | 114 |
| PER260A1 | 60 | 3634752 | 3351627 | 92% | 3137123 | 94% | 3259404 | 97% | 372656 | 361263 | 97% | 346747 | 93% | 352173 | 95% | 145 |
| RO393A1 | 50 | 859962 | 598855 | 70% | 571336 | 95% | 569962 | 95% | 108602 | 101921 | 94% | 93609 | 86% | 97702 | 90% | 138 |

**Supplementary Table 6:** Results from prediction of guide RNA genes within four isolates, representing the three major *Leishmania* populations in Peru and a hybrid *L. braziliensis* x *L. peruviana* isolate.

| Isolate | Column1 | LCA04 | HR78 | HR434 | LC1412 |
|---|---|---|---|---|---|
| Species | | *L. peruviana* | *L. peruviana* | Hybrid | *L. braziliensis* |
| Population | | SUCS | PORCULLA | - | - |
| Number of minicircles | | 80 | 106 | 141 | 117 |
| Number of annotated minicircles | | 65 | 83 | 91 | 93 |
| Percentage of annotated minicircles | | 81% | 78% | 65% | 79% |
| Number of predicted guide RNA genes in minicircles | | 104 | 130 | 138 | 135 |
| Number of predicted guide RNA genes in maxicircle | | 19 | 21 | 19 | 19 |
| Total number of gRNAs | | 123 | 151 | 157 | 154 |
| **Number of guide RNA genes per maxicircle gene (maxicircle-encoded gRNA's are given in brackets):** | | | | | |
| A6 | 5' edited | 5 (2) | 7 (1) | 7 (1) | 6 (1) |
| COIII | 5' edited | 2 | 1 | 2 | 2 |
| CYb | 5' edited | 0 (2) | 1 (2) | 1 (2) | 0 (2) |
| GR3 | pan edited | 4 (3) | 5 (4) | 5 (4) | 6 (4) |
| GR4 | pan edited | 25 (5) | 29 (5) | 32 (6) | 31 (6) |
| ND3 | pan edited | 18 | 24 | 25 | 20 |
| ND7 | 5' edited | 2 (1) | 1 (1) | 3 (1) | 1 (1) |
| ND8 | pan edited | 14 (1) | 19 (2) | 20 (1) | 24 (1) |
| ND9 | pan edited | 20 (4) | 29 (5) | 30 (3) | 31 (3) |
| RPS12 | pan edited | 14 | 14 | 13 | 14 |
| COII | internally edited | 0 | 0 | 0 | 0 |
| MURF2 | 5' edited | 0 (1) | 0 (1) | 0 (1) | 0 (1) |
| **Coverage statistics:** | | | | | |
| initiator gRNAs | | 7 | 9 | 9 | 10 |
| gRNAs with mismatches | | 109 | 133 | 138 | 131 |
| missing gRNAs | | 16 | 10 | 12 | 9 |
| insertions | | 1589 | 1583 | 1588 | 1588 |
| insertions covered | | 1483 | 1537 | 1543 | 1556 |
| insertions not covered | | 106 | 46 | 45 | 32 |
| deletions | | 162 | 162 | 160 | 160 |
| deletions covered | | 137 | 158 | 159 | 160 |
| deletions not covered | | 25 | 4 | 1 | 0 |
| Percentage covered | | 92,52% | 97,13% | 97,37% | 98,17% |

**Supplementary Table 7:** Predicted maxicircle-encoded guide RNA genes. Note that for isolate LCA04 there is no annotation after position 17,206 due to an incomplete maxicircle assembly. Orange bars show guide RNA genes also found in *L. tarentolae.* Yellow bars indicate guide RNA genes unique to *L. braziliensis* .

| strain | start on maxi | end on maxi | strand | gene | start on gene | end on gene | strain | start on maxi | end on maxi | strand | gene | start on gene | end on gene | strain | start on maxi | end on maxi | strand | gene | start on gene | end on gene | strain | start on maxi | end on maxi | strand | gene | start on gene | end on gene |
|---|---|---|---|---|---|---|---|---|---|---|---|---|---|---|---|---|---|---|---|---|---|---|---|---|---|---|---|
| HR78 | 234 | 281 | template | ND9 | 563 | 610 | LCA04 | 133 | 180 | template | ND9 | 563 | 610 | | | | | | | | | | | | | | |
| HR78 | 363 | 410 | coding | ND9 | 205 | 252 | LCA04 | 262 | 309 | coding | ND9 | 205 | 252 | LC1412 | 1246 | 1293 | coding | ND9 | 205 | 252 | HR434 | 1246 | 1293 | coding | ND9 | 205 | 252 |
| HR78 | 405 | 453 | template | GR3 | 142 | 190 | LCA04 | 304 | 352 | template | GR3 | 142 | 190 | LC1412 | 1288 | 1336 | template | GR3 | 142 | 190 | HR434 | 1288 | 1336 | template | GR3 | 142 | 190 |
| | | | | | | | LCA04 | 344 | 384 | template | A6 | 135 | 175 | | | | | | | | | | | | | | |
| HR78 | 557 | 600 | coding | ND9 | 90 | 133 | LCA04 | 456 | 499 | coding | ND9 | 90 | 133 | LC1412 | 1439 | 1482 | coding | ND9 | 90 | 133 | HR434 | 1439 | 1482 | coding | ND9 | 90 | 133 |
| HR78 | 689 | 737 | template | ND7 | 17 | 65 | LCA04 | 588 | 636 | template | ND7 | 17 | 65 | LC1412 | 1571 | 1619 | template | ND7 | 17 | 65 | HR434 | 1571 | 1619 | template | ND7 | 17 | 65 |
| HR78 | 1577 | 1623 | coding | GR4 | 24 | 70 | LCA04 | 1476 | 1522 | coding | GR4 | 24 | 70 | LC1412 | 2459 | 2505 | coding | GR4 | 24 | 70 | HR434 | 2459 | 2505 | coding | GR4 | 24 | 70 |
| HR78 | 2595 | 2645 | template | CYb | 21 | 71 | LCA04 | 2489 | 2539 | template | CYb | 21 | 71 | LC1412 | 3532 | 3582 | template | CYb | 21 | 71 | HR434 | 3532 | 3582 | template | CYb | 21 | 71 |
| HR78 | 8444 | 8495 | coding | ND9 | 304 | 355 | LCA04 | 8336 | 8387 | coding | ND9 | 304 | 355 | LC1412 | 9368 | 9419 | coding | ND9 | 304 | 355 | HR434 | 9368 | 9419 | coding | ND9 | 304 | 355 |
| HR78 | 8590 | 8643 | coding | GR4 | 251 | 304 | LCA04 | 8482 | 8535 | coding | GR4 | 251 | 304 | LC1412 | 9514 | 9567 | coding | GR4 | 251 | 304 | HR434 | 9514 | 9567 | coding | GR4 | 251 | 304 |
| HR78 | 10790 | 10836 | template | GR3 | 66 | 112 | LCA04 | 10682 | 10728 | template | GR3 | 66 | 112 | LC1412 | 11714 | 11760 | template | GR3 | 66 | 112 | HR434 | 11714 | 11760 | template | GR3 | 66 | 112 |
| HR78 | 11117 | 11158 | template | GR4 | 81 | 122 | LCA04 | 11009 | 11050 | template | GR4 | 81 | 122 | LC1412 | 12041 | 12082 | template | GR4 | 81 | 122 | HR434 | 12041 | 12082 | template | GR4 | 81 | 122 |
| HR78 | 11117 | 11158 | template | ND8 | 1 | 42 | LCA04 | 11009 | 11050 | template | ND8 | 1 | 42 | LC1412 | 12041 | 12082 | template | ND8 | 38 | 79 | HR434 | 12041 | 12082 | template | ND8 | 38 | 79 |
| HR78 | 11128 | 11169 | template | GR4 | 248 | 289 | LCA04 | 11020 | 11061 | template | GR4 | 248 | 289 | LC1412 | 12052 | 12093 | template | GR4 | 248 | 289 | HR434 | 12052 | 12093 | template | GR4 | 248 | 289 |
| HR78 | 11586 | 11628 | coding | GR3 | 97 | 139 | LCA04 | 11478 | 11520 | coding | GR3 | 97 | 139 | LC1412 | 12510 | 12552 | coding | GR3 | 97 | 139 | HR434 | 12510 | 12552 | coding | GR3 | 97 | 139 |
| HR78 | 13452 | 13509 | coding | MURF2 | 26 | 83 | LCA04 | 13344 | 13401 | coding | MURF2 | 26 | 83 | LC1412 | 14376 | 14433 | coding | MURF2 | 26 | 83 | HR434 | 14376 | 14433 | coding | MURF2 | 26 | 83 |
| HR78 | 17077 | 17123 | template | CYb | 48 | 94 | LCA04 | 16969 | 17015 | template | CYb | 48 | 94 | LC1412 | 18001 | 18047 | template | CYb | 48 | 94 | HR434 | 18001 | 18047 | template | CYb | 48 | 94 |
| HR78 | 17213 | 17259 | coding | GR4 | 399 | 445 | LCA04 | 17105 | 17151 | coding | GR4 | 399 | 445 | LC1412 | 18137 | 18183 | coding | GR4 | 399 | 445 | HR434 | 18137 | 18183 | coding | GR4 | 399 | 445 |
| HR78 | 17314 | 17359 | template | A6 | 114 | 159 | LCA04 | 17206 | 17251 | template | A6 | 112 | 157 | LC1412 | 18231 | 18271 | template | A6 | 117 | 157 | HR434 | 18231 | 18271 | template | A6 | 117 | 157 |
| HR78 | 18651 | 18695 | coding | ND8 | 381 | 425 | | | | | | | | LC1412 | 19636 | 19680 | coding | GR4 | 254 | 298 | HR434 | 19636 | 19680 | coding | GR4 | 254 | 298 |
| HR78 | 18671 | 18715 | coding | GR3 | 94 | 138 | | | | | | | | LC1412 | 19670 | 19711 | coding | GR3 | 153 | 194 | HR434 | 19670 | 19711 | coding | GR3 | 153 | 194 |
| HR78 | 18674 | 18717 | coding | ND9 | 322 | 365 | | | | | | | | | | | | | | | | | | | | | |

946
